# A novel HERC4-dependent glue degrader targeting STING

**DOI:** 10.1101/2023.02.08.527642

**Authors:** Merve Mutlu, Isabel Schmidt, Andrew I. Morrison, Benedikt Goretzki, Felix Freuler, Damien Begue, Nicolas Pythoud, Erik Ahrne, Sandra Kapps, Susan Roest, Debora Bonenfant, Delphine Jeanpierre, Thi-Thanh-Thao Tran, Rob Maher, Shaojian An, Amandine Rietsch, Florian Nigsch, Andreas Hofmann, John Reece-Hoyes, Christian N. Parker, Danilo Guerini

**Author notes:** Amsterdam UMC location Vrije Universiteit Amsterdam, Molecular Cell Biology & Immunology, Amsterdam institute for Infection and Immunity, De Boelelaan 1117, Amsterdam, The Netherlands.

## Abstract

Stimulator of interferon genes (STING) is a central component of the pathway sensing the presence of cytosolic nucleic acids, having a key role in type I interferon innate immune response. Localized at the endoplasmic reticulum (ER), STING becomes activated by cGAMP, which is generated by the intracellular DNA sensor cyclic GMP-AMP synthase (cGAS). Due to its critical role in physiological function and its ‘ involvement in a variety of diseases, STING has been a notable focus for drug discovery. Recent advances in drug discovery allow the targeting of proteins previously considered “un-druggable” by novel mechanism of actions. Molecular glue degraders are defined as the compounds leading targeted protein degradation (TPD) by creating novel ligase-substrate interactions. Here, we identified AK59 as a novel molecular glue degrader for STING. A genome-wide, CRISPR/Cas9 knockout screen showed that the compound-mediated degradation of STING by AK59 is compromised by the loss of HECT and RLD domain containing E3 ubiquitin protein ligase 4 (HERC4), ubiquitin-like modifier activating enzyme 5 (UBA5) and ubiquitin like modifier activating enzyme 6 (UBA6). While UBA5 and UBA6 could be the auxiliary factors for AK59 activity, our results indicate that HERC4 is the main E3 ligase for the observed degradation mechanism. Validation by individual CRISPR knockouts, co-immunoprecipitations, as well as proximity mediated reporter assays suggested that AK59 functions as a glue degrader by forming a novel interaction between STING and HERC4. Furthermore, our data reveals that AK59 was effective on the most common pathological STING mutations that cause STING-associated vasculopathy with onset in infancy (SAVI), suggesting a potential clinical application of this mechanism. Thus, these findings not only reveal a novel mechanism for compound-induced degradation of STING but also utilize HERC4 as potential E3 ligase that for TPD, enabling novel therapeutic applications.

## Introduction

The cyclic GMP–AMP synthase (cGAS)–stimulator of interferon genes (STING) pathway is an important element of the innate immune system for the detection of, and the response to aberrant intracellular DNA molecules. The pathway is activated by the binding of cGAS protein to cytosolic double-stranded (DNA); this interaction initiates the production of 2′3′ cyclic GMP–AMP (cGAMP) which binds to STING (previously known as TMEM173 or STING1). Located on the endoplasmic reticulum (ER) membrane in its ‘ naive state, STING then dimerizes in the presence of cGAMP; it is translocated from the ER membrane and eventually triggers downstream interferon signaling by a cascade initiated by TANK binding kinase (TBK) phosphorylation ^1^. The presence of double-stranded DNA in the cytoplasm is associated with viral/bacterial infection, retroviral replication and cellular stress (which could be caused by the release of mitochondrial DNA) and so the production of cGAMP, and activation of STING, acts as an innate immune response to such stimuli ^2^.

Aberrant activation of cGAS/STING has been associated with numerous autoimmune diseases ^3^. Ataxia-telangiectasia (AT), a genetic disorder linked to ATM kinase loss of function, as well as Aicardi-Goutieres syndrome (AGS), a relatively rare inflammatory disease, are among the diseases that were linked to cGAS/STING pathway activity ^4,5^. Furthermore, individual point mutations of the STING protein are causative of STING-associated vasculopathy with onset in infancy (SAVI disease) which causes skin lesions, rashes and interstitial lung disease ^6^. The links between the cGAS/STING pathway to multiple autoinflammatory diseases justify efforts to pharmacologically target members of this pathway ^3,7^. As one of the main elements of this pathway, STING has been a prominent target of such efforts ^8,9^. While a straight forward approach to STING inhibition could be targeting the cGAMP binding pocket ^10^, which is the domain responsible for activation, there are many other mechanisms to antagonize a target protein, such as targeting post-translational modifications ^8^.

Ubiquitin signaling is an important <u>p</u>ost<u>t</u>ranslational <u>m</u>odification (PTM) of proteins that regulates numerous signaling pathways in eukaryotic cells ^11^. The main elements of the ubiquitin-proteosome system (UPS) consist of ubiquitin-activating enzymes (E1), ubiquitin-conjugating enzymes (E2) and ubiquitin ligases (E3). As their names suggest, the chain reaction linking ubiquitination of substrate proteins starts with ubiquitin activation via an E1 enzyme, subsequently the ubiquitin is transferred, by an E2 enzyme, from the E1 component to an E3 ligase which links ubiquitin to the target protein. The function of the E3 ligase is to create a scaffold for both the E2 and substrate for transfer of the ubiquitin to the target protein ^12^. There are four major E3 ligase families clustered according to their functional domains and mechanism of action. A major cluster of E3 ligases, which include representatives such as APC, SCF complex or DCX complex, are members of the RING (Really Interesting New Gene)-finger containing family ^13,14^. While RING-finger-containing E3 ligases function as scaffolds mediating close proximity between E2 and the target protein, the HECT-domain-containing E3 ligases interact with ubiquitin directly, utilizing an intermediate step before ubiquitination of target protein ^15^. HERC4 belongs to the HECT-domain containing E3 ligase family and is relatively less well characterized than the RING domain family of E3 ligases. HERC4 expression has been associated with several cancer types including breast cancer, hepatocellular carcinomas and lung tumors ^16–18^. Additionally, only a few target substrates of HERC4 have been identified; such as Smoothened protein (SMO) ^19,20^, Salvador (SAV) protein ^21^, c-Maf ^22^ and progesterone receptor (PGR) ^23^. The structure and the mechanism of action of HERC4 has not yet been fully defined.

Interest in E3 ligases has increased tremendously over the last 20 years due the discovery of compounds that can bring about novel protein interactions with E3 ligases. Molecular glue degraders and proteolysis-targeting chimera (PROTACs) defined as compounds that mediate association of an E3 ligase to a target protein that is usually not a natural substrate of that E3 ligase. This chemically mediated protein-protein interaction (PPI) brought completely new perspective to the field of drug discovery, allowing proteins to be targeted for proteasomal degradation (TPD).

Our study leveraged genome-wide CRISPR/Cas9 knockout screening to enumerate the genes involved in a novel compound-directed STING degradation mechanism. This report describes the identification of the novel compound (AK59-51TB, in short AK59) that inhibits the cGAS/STING pathway via targeted proteasomal degradation of STING. A genome-wide CRISPR-Cas9 screen was conducted to identify genes inhibiting the compound-mediated STING degradation. Subsequently, the function of AK59 was shown to be dependent on HERC4, UBA5 and UBA6, each of which reduces AK59 activity when individually knocked out. Moreover, immunoprecipitation assays and a novel PPI assay revealed that the interaction between STING and HERC4 only occurs in the presence of the compound. These results identify AK59 as a glue degrader of STING and demonstrate a novel regulatory mechanism of the cGAS/STING pathway, unveiling the potential of HERC4 as an E3 ligase that has not previously been exploited for known TPD mechanisms.

## Results

### AK59 is a selective inhibitor of STING

Due to its ‘ significance in various autoimmune diseases and cancer, the regulation of the cGAS/STING pathway is of interest to pharmaceutical companies and society^3^. STING is one of the main elements of this pathway and a key regulator of interferon regulatory factors (IRFs), thus regulation of STING expression may have a high potential therapeutic value in the field of immunology ^9^. Due to this potential therapeutic application, a portion of the Novartis compound collection was screened to identify compounds that inhibit the type 1 interferon response upon expose to exogenous DNA (data not shown). THP1 human monocytic cells were chosen as a suitable cell model with detectable and relevant cGAS/STING pathway activity. Among the compounds screened, AK59 showed significant inhibitory effect on STING activity. Western blot analysis of THP1 cells showed that 16 hours of compound incubation caused a drastic decrease of STING protein levels (Figure 1A). While AK59 showed significant decrease in STING protein levels, QK50-66NB (in short QK50) treatment, a close structural analog of AK59, did not affect STING levels in treated THP1 cells (Figure 1A), highlighting the specificity of AK59. To demonstrate that STING degradation translated to downstream inhibition of the pathway, commercially available Dual-THP1-Cas9 cells (that have a Lucia luciferase reporter linked to IRF responsive promoter) were used to monitor compound effects on activation of the interferon response pathway. Upon stimulation with 30 µM of cGAMP, Dual-THP1-Cas9 cells showed an increased luminescence signal that was significantly inhibited by treatment with 10 µM AK59 (Figure 1B). Dual-THP1-Cas9 cells in which STING had been knocked out ^24^ were used as a negative control to show the specificity of the pathway activation via cGAMP stimulation. Additionally, dose response of these two related compounds showed that QK50 treatment of cGAMP stimulated Dual-THP1-Cas9 cells did not show the same decrease in IRF pathway signaling (Supplementary Figure S1A). Both western blot as well as reporter assay results showed that AK59 was specifically inhibiting STING-mediated activation of the IRF pathway.

**Figure 1.**
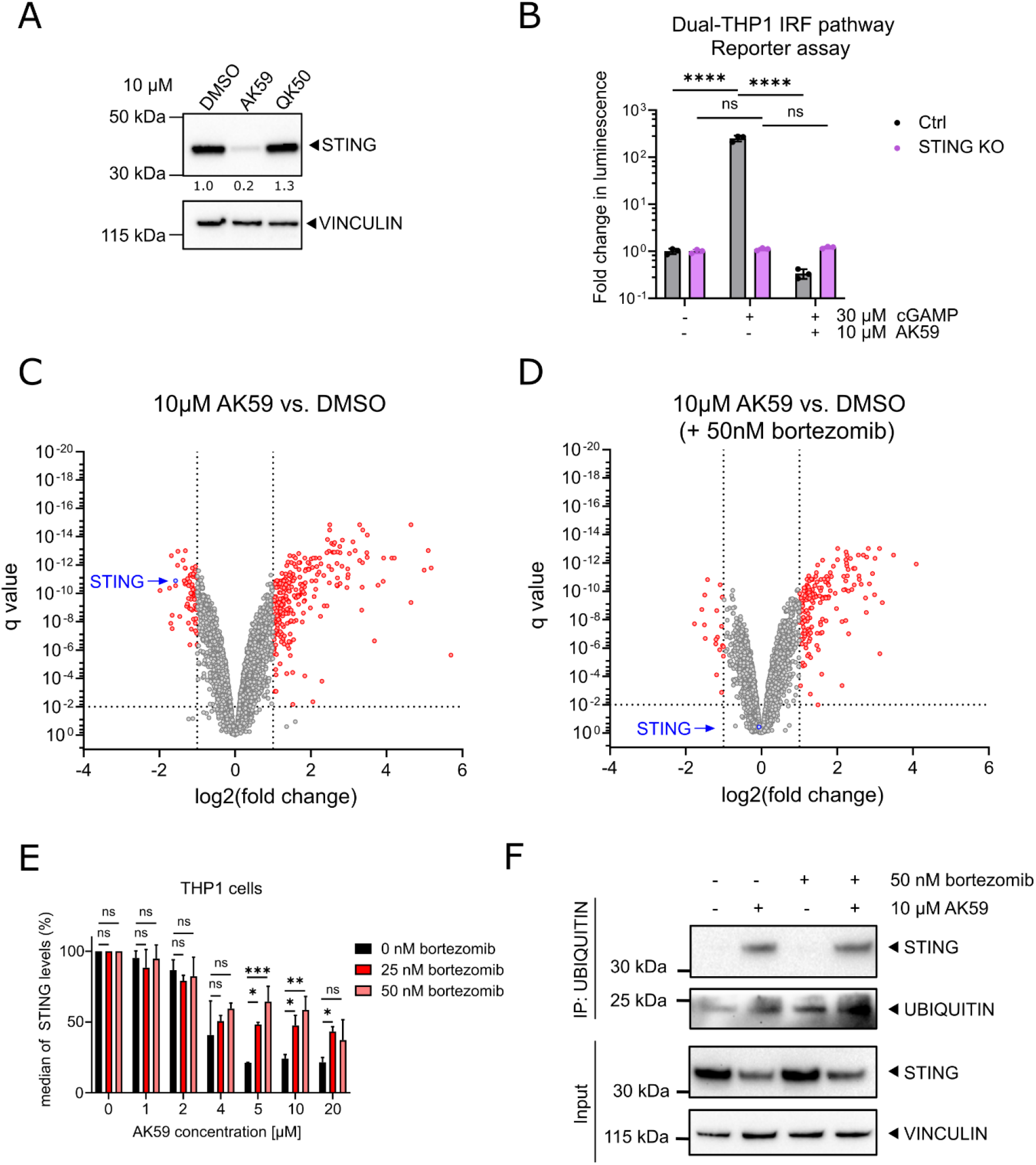
AK59 decreases STING protein on THP1 cells through proteasomal degradation. **(a)** Western blot showing STING protein levels in THP1 cells treated with either AK59 or QK50 for 16 hours. VINCULIN was used as a loading control. The STING expression quantified by first normalizing to its respective loading control and ratio to DMSO control. **(b)** IRF pathway reporter assay on wild type or STING knockout Dual-THP1-Cas9 cells. Cells were stimulated with 30 µM cGAMP 3 hours prior to 16 hours 10 µM AK59 incubation. Luminescence reads were normalized to the control (unstimulated) sample for each cell line. Statistical significance was calculated using two-way ANOVA followed by Šidák ‘s correction. Data plotted as mean ± SD of three individual biological replicates in Graphpad Prism (Version 9). **(c-d)** Volcano plots of results from the proteomics analysis. THP1 cells were for 16 hours with 10 µM AK59 compared to control DMSO treated samples. Significantly altered protein abundances are shown with a log2 fold change < - 1 or > 1 and a q value of less than 0.01. **(e)** STING expression of THP1 cells treated with AK59 in the presence of 25nM or 50nM prior bortezomib treatment. STING expression was detected with FACS analysis on fixed cells. Bortezomib treatment was 1 hour prior to AK59 treatments. Each treatment group normalized to its individual vehicle control and represented in percentage. Three biological replicates were plotted as mean ± SD using in Graphpad Prism (Version 9). Statistical significance was calculated using two-way ANOVA followed by Šidák ‘s correction. **(f)** Ubiquitin pulldown followed by western blot on THP1 cells treated either 10 µM AK59 or 50 nM bortezomib or both. VINCULIN was used as a loading and pulldown control.

To further characterize the effect of AK59 on THP1 cells, an unbiased proteomics approach used. THP1 cells treated with either DMSO (vehicle), 10 µM AK59 or 10 µM QK50 for 16 hours and tryptic digests of total cell lysates were analyzed by LC-MS/MS (Supplementary Figure S1B). The treatment conditions were compared to the vehicle control group and fold changes were calculated accordingly. The proteomics analysis revealed significantly downregulated proteins with a < -1 Log2 fold decrease and a false discovery rate p-value (q value) of less than 0.01 (Figure 1C, Supplementary Table S1). Protein expression levels of STING was found to be significantly downregulated upon AK59 treatment compared to vehicle control. Furthermore, STING expression levels remained unaltered in the presence of non-functional analog of AK59, QK50 (Supplementary Figure S1C-D).

UPS machinery an important protein regulatory element in eukaryotic cells that plays a critical role in numerous cellular pathways ^25^. To understand whether change of the protein levels of STING is due to proteasomal degradation, bortezomib was added prior to the compound treatment. Changes of the proteomics profile upon 50nM bortezomib addition showed that the downregulation on STING protein by AK59 were rescued (Figure 1D). To validate this finding, proteasomal degradation was inhibited using two different proteasome inhibitors (bortezomib and MG132). In addition, a neddylation inhibitor (MLN4924) was tested on AK59 treated THP1 cells (Supplementary Figure S1E). Proteasomal inhibition or neddylation inhibition was initiated 1-2 hours prior to compound treatment and maintained during the 16 hours of AK59 incubation. Interestingly, neddylation inhibition did not rescued AK59-mediated STING downregulation. While the proteasomal inhibitors were highly toxic to the cells, bortezomib showed the best rescue of the compound mediated STING degradation. Additionally, two concentrations of bortezomib treatment (25 and 50 nM) were selected as having the largest effect on proteasomal inhibition without causing excessive cell death when combined with various doses of AK59. The STING protein levels in THP1 cells that measured by FACS (Figure 1E) showed that proteasomal inhibition via bortezomib rescues compound-derived STING degradation to a certain extend in a dose dependent manner. Furthermore, ubiquitination pulldown assays showed STING ubiquitination upon AK59 (Figure 1F). These findings corroborated an inhibitory effect of AK59 on STING and therefore cGAS/STING signaling, via induced UPS.

### Lysine 150 of STING is essential for AK59-mediated STING degradation

To dissect the functional domain(s) of STING essential for novel AK59-mediated degradation, GFP tagged STING expression vectors were constructed and expressed in HEK293T cells (which do not express endogenous STING). The expression plasmids constructed were focused on the cytosolic domain of STING to avoid the complexity the transmembrane domains might introduce in folding or localization. Several constructs that contain the cytosolic domain of STING differing on the connector helix domain were created (Supplementary Figure S2). STING degradation was tracked in live cells by following GFP fluorescence after 10 µM AK59 treatment. STING expression on STING^141-341^-GFP (construct #1) expressing HEK293T cells was significantly reduced while the shorter STING^155-341^-GFP(construct #4) expressing HEK293T cells seemed not to be influenced by compound treatment (Figure 2A). This indicated that the connector helix domain of STING protein (141-155aa) appears to be involved in the function of AK59. To understand whether this domain is important in the binding or the ubiquitination of the STING protein, lysine residues at the connector helix domain were further investigated. The K150 lysine residue, located within the connector helix domain, has been linked to PTMs such as ubiquitination of STING and its degradation ^26,27^. In order dissect the two functions of AK59 on STING (interaction vs. ubiquitination), mutation of K150 to arginine (K150R) on non-GFP tagged STING^141-341^ construct (STING^141-341_K150R^, construct #6)) was introduced. Together with non-GFP tagged STING^141-341^(construct #5) and STING^155-341^(construct #7), STING^141-341_K150R^ construct transiently expressed in HEK293T for side-by-side comparison. Ubiquitin pulldown followed by western blot on these cell lines showed that ubiquitination levels remained constant in either STING^155-341^ or STING^141-341_K150R^ expressing HEK293T, whereas ubiquitination of STING has been increased in STING^141-341^ expressing HEK293T cells (Figure 2B). These data indicated that K150 residue of STING is essential for AK59-mediated ubiquitination and therefore degradation of the protein.

**Figure 2.**
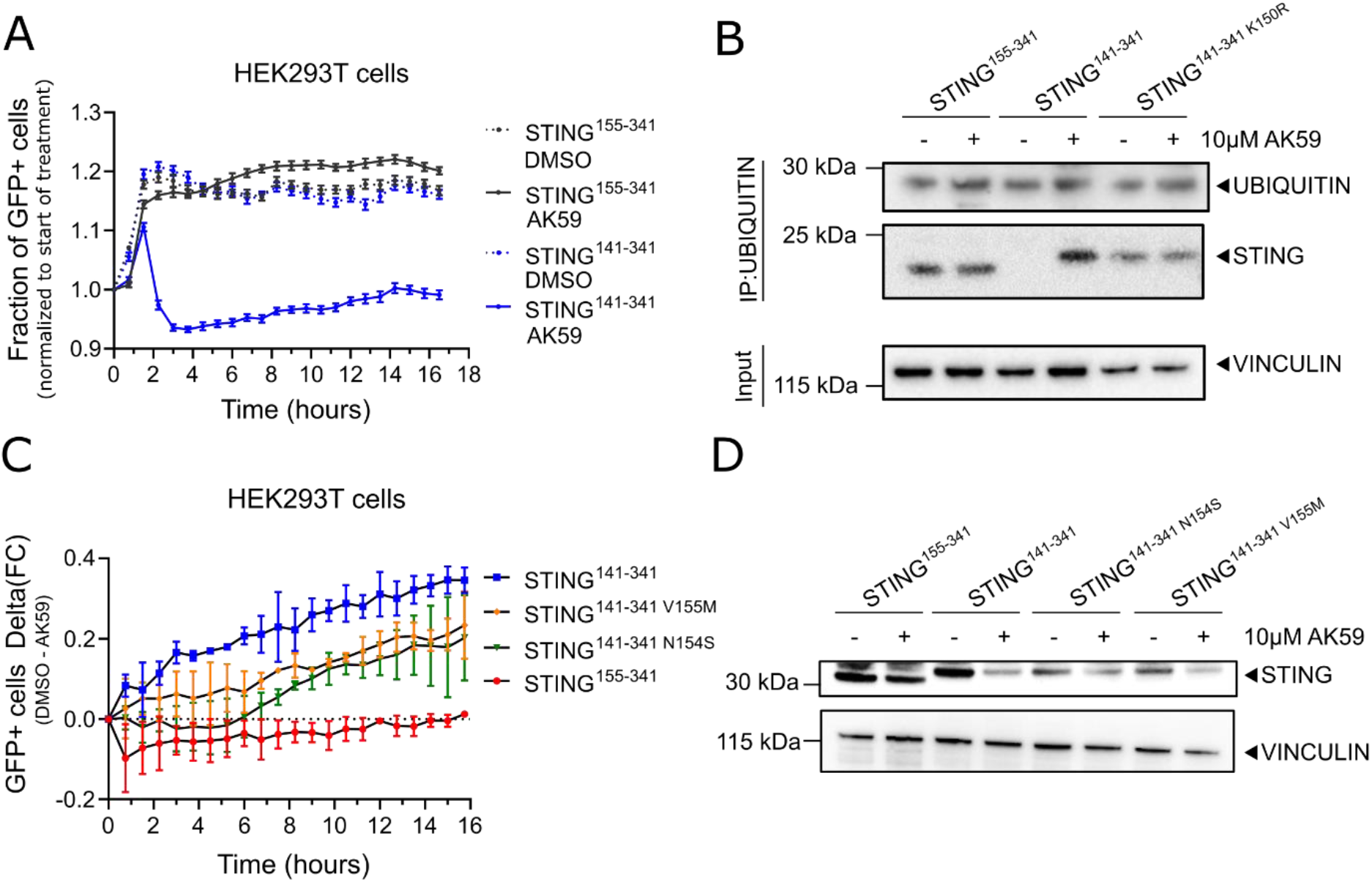
K150 is essential for STING ubiquitination upon AK59 treatment. **(a)** Live GFP positive cell tracking of indicated STING^141-341^-GFP or STING^155-341^-GFP constructs transfected HEK293T cells upon 10 µM AK59 treatment. Data points were normalized to the start of the treatment. For each data point 16 pictures/well was taken. Data from biological replicates were plotted as mean ± SD using in Graphpad Prism (Version 9). **(b)** Ubiquitin pulldown followed by western blot on HEK293T cells transfected with the indicated STING expression constructs and treated either 10 µM AK59 or vehicle control. VINCULIN was used as a loading and pulldown control. **(c)** Live cell tracking of indicated STING-GFP constructs that harbors SAVI mutations transfected HEK293T cells after 10 µM AK59 treatment. Data points were normalized to the start of the treatment. For each data point 16 pictures/well were taken. Data from biological replicates were plotted as mean ± SD using in Graphpad Prism (Version 9). **(d)** Western blot representing the changes in STING protein levels upon AK59 treatment in SAVI mutations harboring STING-GFP expressing HEK293T cells. Indicated constructs were transiently expressed on the cells. VINCULIN used as a loading control.

To investigate whether AK59 functions also on pathological STING variants, cGAS/STING pathway associated diseases searched in the literature. STING-associated vasculopathy with onset in infancy (SAVI) is rare genetic autoimmune disorder caused by single point mutations on STING protein ^28–31^. The point mutations N154S and V155M are present in the cytosolic domain of STING and are the most commonly detected causative mutations in SAVI patients ^32^. In order to observe the effect of AK59 on SAVI mutant STING proteins, these two mutations were introduced into the STING^141-341^-GFP constructs (constructs #1-4, Supplementary Figure S2). After transient transfection of the wild type and mutant STING constructs, GFP signal in live cells were tracked in vehicle treated or 10 µM AK59 treated cells. Tracking of live cells revealed that, similar to STING^141-341^-GFP construct, both STING variants were partially degraded in the presence of AK59 treatment (Figure 2C and Figure 2D). Our data indicate that AK59 is still functional against SAVI mutants to a certain extent, and this paves the way for future translational implications of AK59.

### AK59 function relies on HERC4, UBA5 and UBA6

While identifying the necessary domains and lysine ubiquitination ‘s for AK59-mediated STING degradation, the mechanism of action of the compound remained unknown. To investigate the mechanism of action of AK59, a pooled, genome-wide, CRISPR-Cas9, knockout screen was conducted ^33^. The aim was to identify genes that when knocked out, significantly abrogate the effect of AK59 in reducing the levels of STING protein in THP1-Cas9 cells.

A FACS-based assay to monitor the levels of STING protein was developed and used as a phenotypic readout for the CRISPR-Cas9 screen. Compound concentration and incubation time were optimized according to the median fold-difference on STING staining (Figure 3A). This identified 10 µM AK59 and a 16-hour incubation as giving a distinctive 5-to-6-fold separation in the STING protein levels in our FACS assay. After establishing the FACS-based STING assay, the CRISPR/Cas9 genome-wide screen was conducted (Figure 3B). Screening subjected cells to either DMSO (vehicle) control or 10 µM of AK59 for 16 hours and cells were sorted according to their STING expression. “high” and “low” STING groups were assigned as the higher and lower 25% quartile of the sorted cells. Enriched and depleted sgRNAs in the comparison of high vs. low STING protein levels following treatment, as well as the control group, were presented as a plot displaying RSA p-value (significance of the hits) for guides targeting each gene as well as the magnitude of this effect compared to the other genes (Q3 values) (Figure 3C, Supplementary Figure S3A-C, Supplementary Table S2). To distinguish the genes that are significantly enriched in AK59 high vs. low STING comparison and stable in DMSO high vs. low comparison, the Euclidean distance between the two sets of (Q, RSA) coordinates was calculated (Supplementary Table S2). By ranking the distances, candidate hits that are altered only in the AK59 low vs. high comparison identified. Figure 3D represents the ranked order of the genes, where their knockout resulted in higher STING protein expression in the AK59 treatment group. Among these highly ranked genes, HERC4, UBA5 and UBA6 attracted our attention. HERC4 is a HECT domain containing E3 ligase, however few of its targets have been identified and reported in the literature ^19–23^. UBA5 and UBA6 are ubiquitin-like modifier activating E1 proteins. Individual sgRNAs targeting these three genes show that most of them indeed enriched in the AK59 treatment group with high STING expression compared to the control group (Figure 3E). Additionally, STRING analysis (Version 11.5) ^34^ revealed that interactions between UBA5, UBA6 and HERC4 had been reported in *Homo sapiens*. Importantly, none of these candidates have not been previously linked to STING (Figure 3F). These results increased our confidence on the mechanism of AK59 on STING regulation is through UPS involving HERC4, UBA5 and UBA6 as regulatory factors.

**Figure 3.**
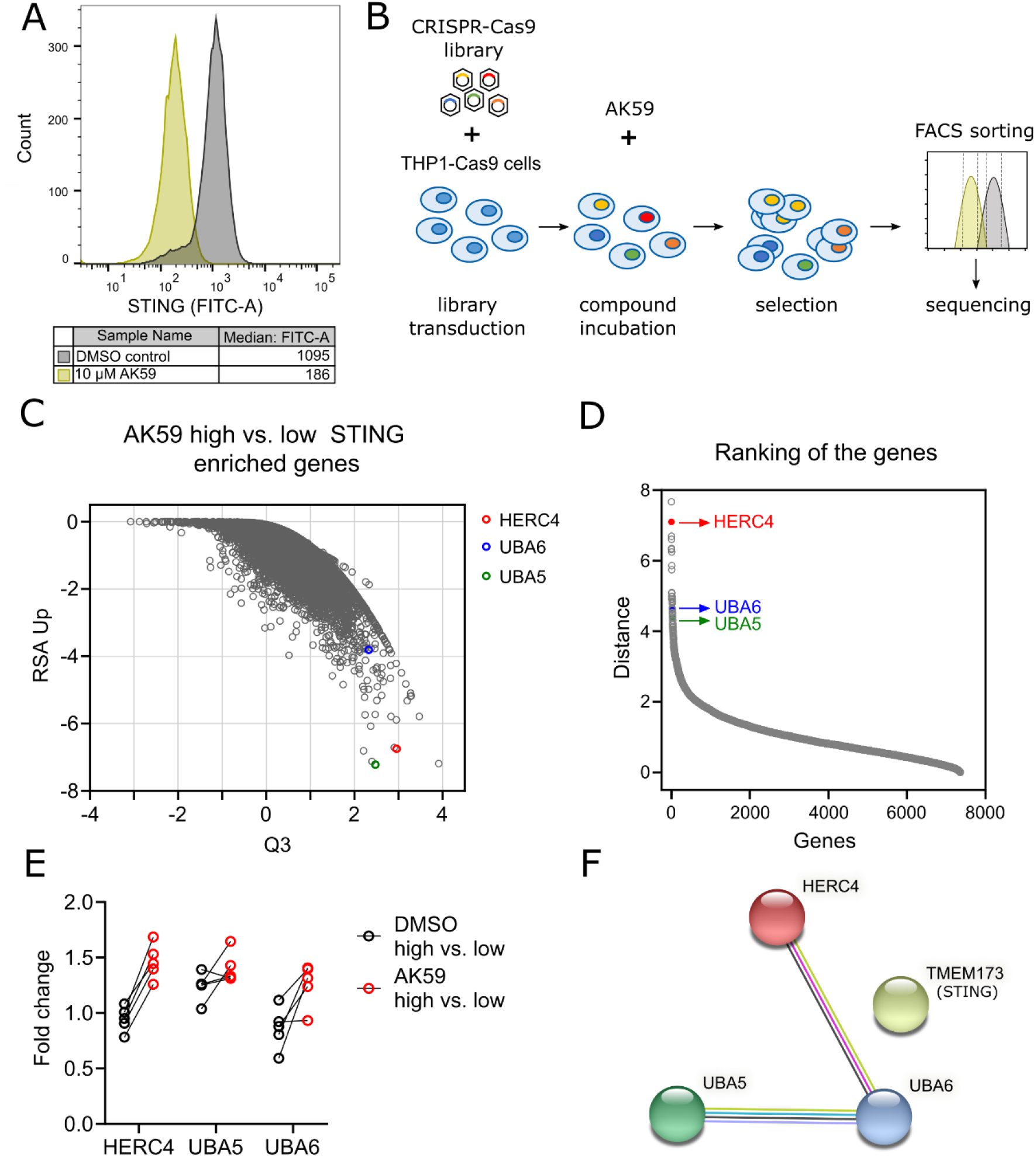
CRISPR/Cas9 genome-wide screening unveiled genes responsible in AK59 activity on STING expression. **(a)** FACS analysis of THP1-Cas9 cells that are treated either with vehicle (green) or 10 µM AK59 (black). Experiment treated at least in three biological replicates and representative FACS reads plotted using FlowJo (Version 10.6.1). **(b)** Schematic representation of the layout of CRISPR/Cas9 genome-wide knockout screen. **(c)** Comparison of RSA Up values with Q3 values from CRISPR/Cas9 screen of 10 µM treated THP1 cells represented in a dot plot. The comparison represented in the blot is high STING expression to low STING expression in the AK59 treatment group. Each dot represents a gene from the CRISPR/Cas9 library. **(d)** Ranking of genes that has enriched sgRNAs in the high STING expression group compared to low. Ranking distance calculated by the change of RSA and Q values between treatment groups. Each gene represented with a dot on the graph. **(e)** Individual fold changes of each sgRNA targeting HERC4, UBA5 and UBA6 in the high vs. low STING expression comparison groups in the compound treated and untreated samples. **(f)** STRING analysis representing the interaction between HERC4, UBA5, UBA6 and STING (STRING version 11.5) ^34^.

To validate the CRISPR genome-wide screen results, individual knockout cell lines constructed. Two best-performing sgRNAs per candidate from the genome library selected and introduced to THP1-Cas9 cells with lentiviral particles. After the third passaging of the cells following lentiviral transduction, cells were collected to check the CRISPR editing efficiency. Modification-rate on the cut-site assessed by PCR followed by tracking of indels by decomposition (TIDE) analysis ^35^ (Figure 4A, 4D and 4G). To show that the modifications on the cut sites on each gene resulted in depletion (or decrease) in protein levels these were assessed (Figure 4B, 4E, 4H and Supplementary Figure S4). ACTIN or VINCULIN proteins were used as a loading control. After confirming the loss-of-function (LoF) of each gene, the FACS based assay monitoring STING protein levels was used to further validate our screen results. HERC4, UBA5 and UBA6 sgRNA transduced THP1-Cas9 lines showed higher levels of STING protein in the presence of 10 µM AK59 treatment compared to control (Ctrl) sgRNA transduced THP1-Cas9 cells. While AK59 treatment resulted in up to 70% decreased of STING protein in the Ctrl cell line, in the HERC4, UBA5 and UBA6 LoF lines this was decrease to 50-60% (Figure 4C, 4F, 4I and Supplementary Figure S4). Furthermore, HERC4 as the second top rated hit in the CRISPR screen showed the highest rescue phenotype of AK59 treatment regarding STING expression (Figure 4J). All these data indicated that we could replicate the results of the genome-wide screen with individual LoF lines and that HERC4, UBA5 and UBA6 loss interrupts the effect of AK59 on reduction of STING protein levels.

**Figure 4.**
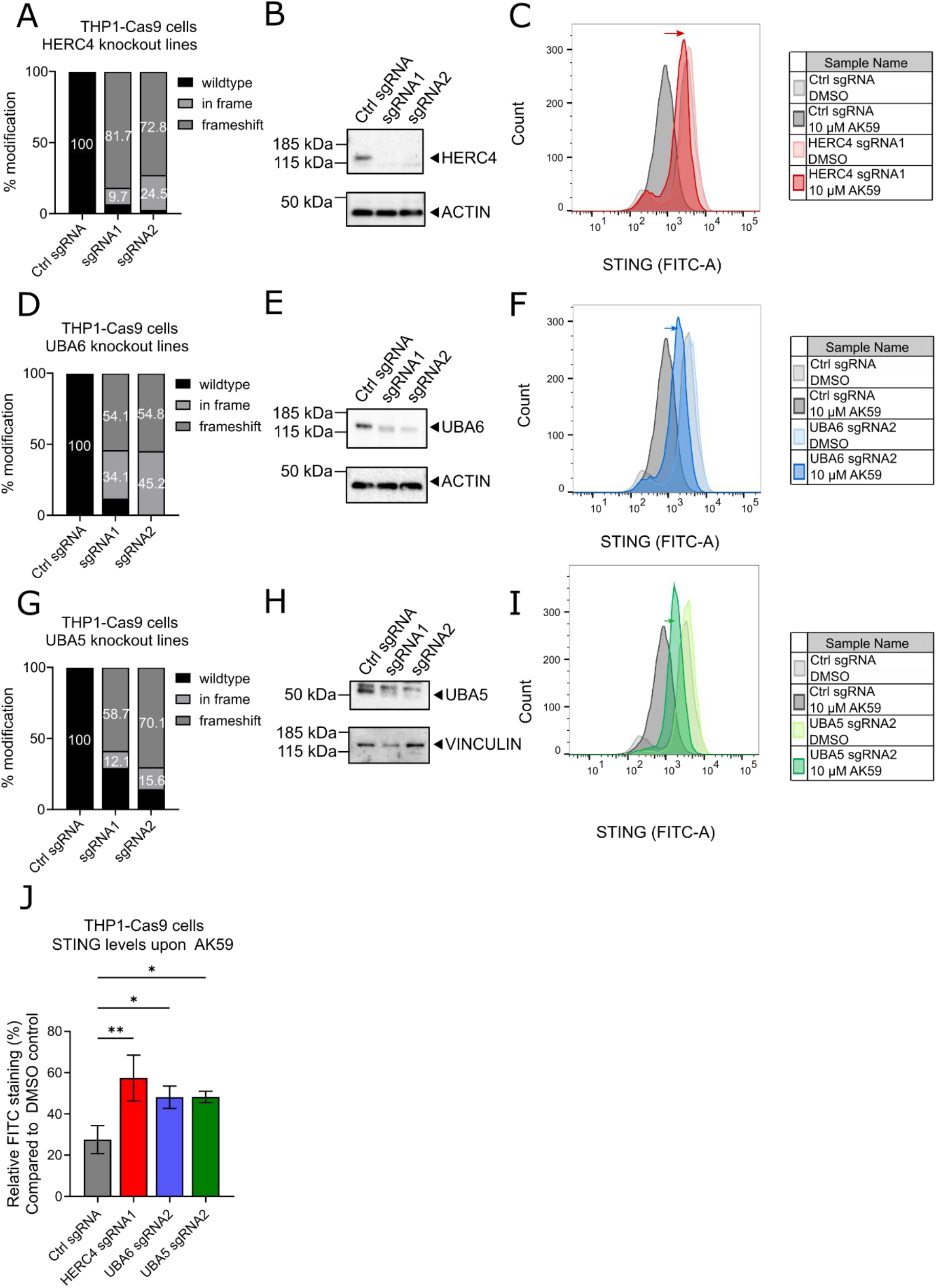
HERC4, UBA5 and UBA6 were validated as genes responsible for AK59 activity on STING levels. **(a)** TIDE analysis of either Ctrl sgRNA, HERC4 sgRNA1 or HERC4 sgRNA2 transduced THP1-Cas9 cells. Modification rates were checked at the third passage after the initial transduction. Undefined sequencing reads were excluded. **(b)** Western blot to detect HERC4 protein levels of either Ctrl sgRNA, HERC4 sgRNA1 or HERC4 sgRNA2 transduced THP1 cells. Proteins from each cell line collected at the third passage after the initial transduction. **(c)** FACS analysis of STING expression on Ctrl sgRNA or HERC4 sgRNA1 transduces THP1-Cas9 cells that were treated either with vehicle (gray and pink) or 10 µM AK59 (black and red). The effect of the knockout indicated with a red arrow. Experiment repeated at least in three biological replicates and representative FACS reads plotted using FlowJo (Version 10.6.1). **(d)** TIDE analysis from either Ctrl sgRNA, UBA6 sgRNA1 or UBA6 sgRNA2 transduced THP1-Cas9 cells. Modification rates were check at the third passage after the initial transduction. Undefined sequencing reads were excluded. **(e)** Western blot to detect UBA6 protein levels on either Ctrl sgRNA, UBA6 sgRNA1 or UBA6 sgRNA2 transduced THP1-Cas9 cells. Proteins from each cell line collected at the third passage after the initial transduction. **(f)** FACS analysis of STING expression on Ctrl sgRNA or UBA6 sgRNA2 transduces THP1 cells that are treated either with vehicle (gray and light blue) or 10 µM AK59 (black and dark blue). The effect of the knockout indicated with a blue arrow. Experiment repeated at least in three biological replicates and representative FACS reads plotted using FlowJo (Version 10.6.1). **(g)** TIDE analysis from either Ctrl sgRNA, UBA5 sgRNA1 or UBA5 sgRNA2 transduced THP1-Cas9 cells. Modification rates were check at the third passage after the initial transduction. Undefined sequencing reads were excluded. **(h)** Western blot to detect UBA5 protein levels on either Ctrl sgRNA, UBA5 sgRNA1 or UBA5 sgRNA2 transduced THP1-Cas9 cells. Proteins from each cell line collected at the third passage after the initial transduction. **(i)** FACS analysis of STING expression on Ctrl sgRNA or UBA5 sgRNA2 transduces THP1-Cas9 cells that were treated either with vehicle (gray and light green) or 10 µM AK59 (black and dark green). The effect of the knockout indicated with a green arrow. Experiment repeated at least in three biological replicates and representative FACS reads plotted using FlowJo (Version 10.6.1). **(j)** Relative STING expressions in indicated THP1-Cas9 cell lines. Data is normalized vehicle treated group and three biological replicates were plotted as mean ± SD using in Graphpad Prism (Version 9). Statistical significance was calculated using one-way ANOVA followed by Dunnett ‘s multiple comparison test.

### AK59 act as a glue degrader modulating novel interaction between STING and HERC4

UBA5 and UBA6 are E1 enzymes suggesting that they may act as accessory elements for AK59-mediated STING degradation. HERC4 was one of the top hits from the CRISPR genome-wide screen and as an E3 ligase became prioritized as having an important role in AK59 function. Furthermore, as HERC4 is an E3 ligase this offered the hypothesis that AK59 is acting as a molecular glue degrader creating a novel interaction between HERC4 and STING, allowing STING to be ubiquitinated and degraded. To test this hypothesis, we first wanted to make sure not only compound-mediated STING degradation, but also cGAS/STING pathway activity was reversed by HERC4 LoF. It was previously demonstrated that the inhibitory effect of AK59 on STING protein was translated to downstream the IRF pathway (Figure 1B). To link HERC4 to AK59 functioning as a mechanism of action, HERC4 sgRNA was introduced to the Dual-THP1-Cas9 ^24^ cell line to create HERC4 LoF. Validation of LoF was shown both by TIDE analysis on the cut site as well as western blot on protein level (Supplementary Figure S5A and S5B). As expected, luminescence due to cGAMP stimulation, indicating activation of the IRF pathway, gradually decreased with increasing concentrations of AK59 in the Ctrl cell line (Supplementary Figure S5C). In contrast to the control cell line, the HERC4 knockout Dual-THP1-Cas9 cell line showed higher luciferase activity in the presence of AK59 indicating that the compound ‘s inhibitory activity on IRF pathway is compromised in the absence of HERC4. Thus, absence of HERC4 interrupted the function of AK59 on STING protein levels and downstream on activation of the IRF pathway.

To observe the changes in the inhibition of cytosolic STING via AK59 in the absence of HERC4, the link between HERC4 and the ubiquitination of STING through AK59 were investigated. First, HERC4 was knocked out in HEK293-JumpIN-Cas9 cell line to test the HERC4 dependent AK59 activity on various STING constructs. Knockout efficiency was shown both by TIDE analysis and western blot (Supplementary Figure S5D and S5E). Furthermore, LoF cell lines were used to transiently expressed GFP-fused cytosolic domain for STING (Supplementary Figure S2). GFP signal tracking on live HERC4 LoF cells that transfected with STING-GFP constructs showed that HERC4 LoF ablates the differences in the degradation of different STING constructs (Figure 5A). In other words, degradation of cytosolic STING via AK59 has been blocked in the absence of HERC4. These results solidified the link HERC4, an E3 ligase, to compound-mediated STING degradation suggesting that AK59 could be functioning as a glue degrader. To further support this data, protein levels known substrates of HERC4 such as SMO and PGR^19,20,23^ were measured in the presence of AK59 (Figure 5B). Both in the wildtype THP1-Cas9 cells as well as STING KO THP1-Cas9 cells, protein levels of SMO as well as PGR remained unchanged or even stabilized upon treatment suggesting that AK59 function on HERC4 is STING-specific. This data is supported by the ubiquitin pulldown in STING^141-341^ expressing HEK293T cells. As expected transiently expressed STING construct shows increased ubiquitination levels in the presence of the AK59 while ubiquitination of SMO remained stable (Figure 5C). Additionally, HERC4, as a HECT domain containing E3 ligase shown to be also degraded in the presence of AK59 (Figure 5B and 5C) which could be due to its ‘ self-ubiquitination during the transfer of the ubiquitin domain from E2 enzyme to substrate.

**Figure 5.**
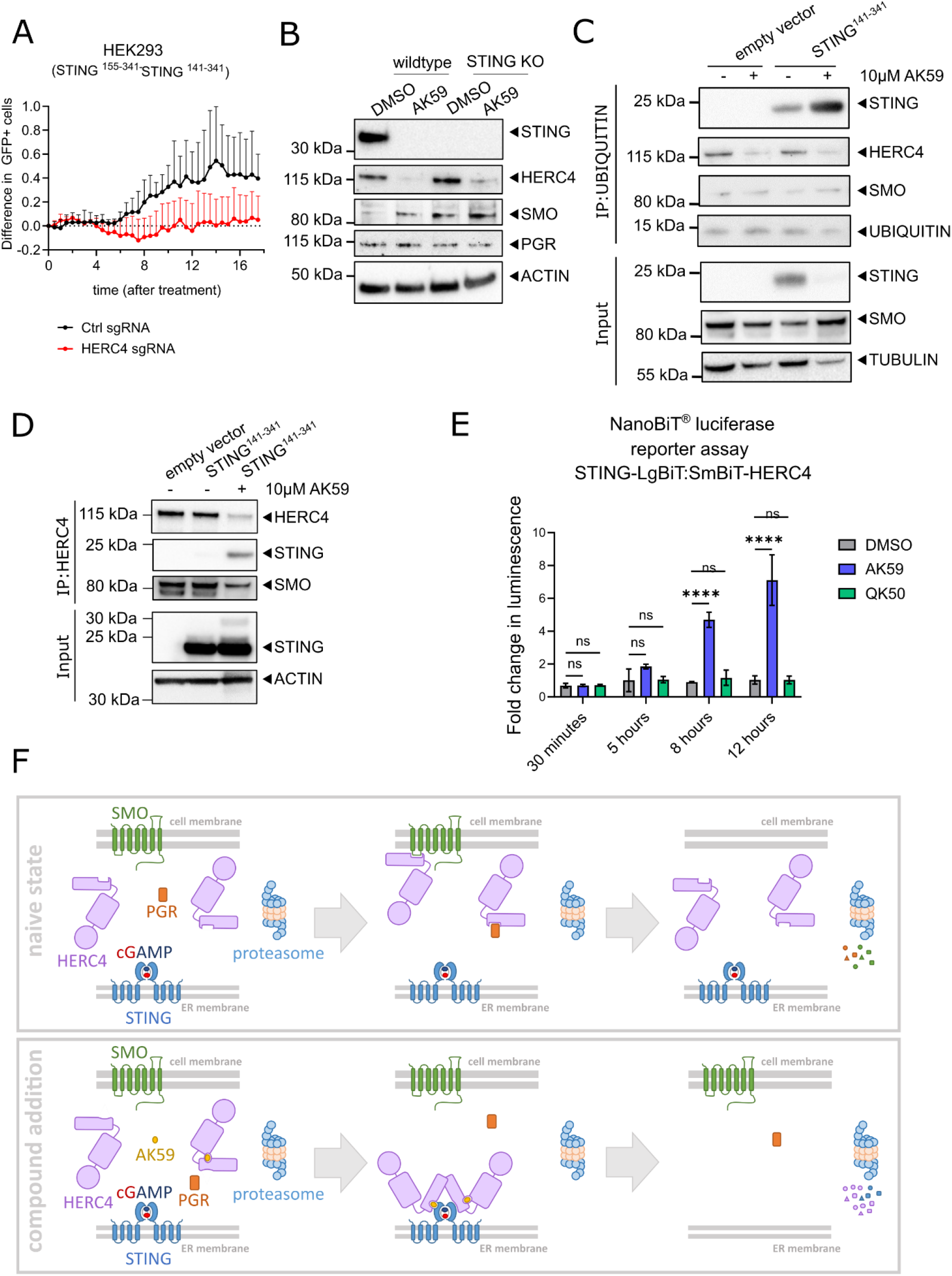
AK59 acts as a glue degrader targeting STING. **(a)** Live cell tracking of wildtype or HERC4 knockout HEK293-JumpIN-Cas9 cells after 10 µM AK59 treatment. Data points were normalized to the moment of AK59 treatment and to their corresponding vehicle control group. For each data point 16 pictures/well were taken. Data from biological replicates were plotted as mean ± SD using in Graphpad Prism (Version 9). **(b)** Western blot showing STING, HERC4, SMO and PGR protein levels in THP1 cells treated with either AK59 or vehicle control for 16 hours. ACTIN was used as a loading control. **(c)** Ubiquitin pulldown followed by western blot on HEK293T cells transfected with the indicated STING expression constructs and treated either 10 µM AK59 or vehicle control. TUBULIN was used as a loading and pulldown control. **(d)** Co-immunoprecipitation followed by western blot on HEK293T cells transiently expressing indicated STING expression constructs and treated either with 10 µM AK59 or vehicle control. ACTIN was used as a loading and pulldown control. **(e)** Nanobit complementation assay on STING-LgBiT and SmBiT-HERC4 expressing HEK293T cells treated with either 10 µM AK59, 10 µM QK50 or vehicle control. Luminescence detected at the indicated time points after the start of the compound treatment. Luminescence reads were normalized to corresponding DMSO control. Three biological replicates were plotted as mean ± SD using in Graphpad Prism (Version 9). Statistical significance was calculated using two-way ANOVA followed by Šidák ‘s correction. **(f)** Graphical representation of naive and compound added state of the cells and the suggested new protein-protein interactions in the presence of AK59.

For AK59 to act as a glue degrader, physical interaction between E3 ligase and its ‘ novel substrate needed to be shown. In order to demonstrate that, experiments were conducted to investigate compound-dependent PPI between HERC4 and STING protein. Co-immunoprecipitation of STING followed by western blot was performed on transiently STING^141-341^ expressing HEK293T cells. HEK293T cells were treated with DMSO or AK59, 48 hours after transfection with either empty vector or STING^141-341^ construct. Co-immunoprecipitation followed by western blots showed that only in the presence of AK59 was STING pulled down together with HERC4 (Figure 5D, 3^rd^ lane). Consecutively, HERC4-interacting SMO protein levels are decreased supporting that AK59 alters substrate recognition of HERC4. DMSO treated STING^141-341^ expressing HEK293T lysate (Figure 5D, 2^nd^ lane) did not show any band on the STING blot and showed no change in SMO levels indicating that the interaction between HERC4 and STING is a novel interaction and dependents on the presence of the compound. Additionally, STING^141-341^, STING^155-341^ and STING^141-341 K150R^ expressing HEK293T cells were used to check the interaction of STING with HERC4. Interestingly, all the STING forms were found to still be interacting shown by co-immunoprecipitation upon compound incubation, which is further validating that connector helix sequence and the K150 residue of STING is only essential for the ubiquitination of the substrate but not for the interaction of the tripartite complex formed in the presence of AK59 (Supplementary Figure S6A).

To support the pulldown data indicating the AK59-mediated novel interaction of HERC4 and STING, by demonstrating compound-dependent proximity, a NanoBiT^®^ complementation assay (Promega) was developed. The Nanobit complementation assay consists of a small (SmBiT) and large (LgBiT) piece of the Nanoluciferase protein, which have low affinity for each other. Only in the presence of a direct interaction of the conjugated proteins domains, can the SmBiT and LgBiT form a functional Nanoluc and produce a luminescence signal ^36^. Expression constructs where the STING 141-341aa cytosolic domain was C-terminally conjugated with the LgBiT domain and full length HERC4, N-terminally linked to the SmBiT were created (Supplementary Figure S6B). These constructs were then transiently transfected into HEK293T cells and expression of the constructs were validated with western blot (Supplementary Figure S6B, below). 48 hours post-transfection; cells were treated with either DMSO or AK59. Time course (Figure 5E) as well as the dose-dependent (Supplementary Figure S6D) measurements showed that LgBiT and SmBiT interaction and therefore the nano luminescence signal is detected only in the presence of AK59. QK50 a non-functional analog of AK59 (Supplementary Figure S1A), was used as a negative control. Furthermore, cGAMP supplementation did not compete with AK59 (Supplementary S6B) indicating that cGAMP is not competing with AK59 or in other words AK59 is not binding to the cGAMP binding site. Specificity of the luminescence signal, only in the presence of AK59 but not in DMSO or QK50 treated groups supports the hypothesis that AK59 induces a novel PPI between STING and HERC4, consistent with the pulldown results.

These results are consistent with activity of AK59 as a novel STING glue degrader. Overall, these findings shows that AK59 is a novel degrader and causes a novel interaction between STING and HERC4. AK59 interacts with HERC4, a HECT domain containing E3 ligase, causing a novel interaction with STING, which results in proteasomal degradation of STING protein. This interaction results in not only the degradation of STING but also HERC4 leading to stabilization of known substrates of HERC4 (Figure 5F). Furthermore, AK59-mediated ubiquitination of STING K150 and subsequent STING degradation results in the inhibition of IRF pathway activation. Additionally, these results showed that AK59 not only degrades wild-type STING but also SAVI mutant STING proteins, with implications for translational applications of the AK59 mechanism of action for chronic STING activation.

## Discussion

This study identified the mechanism of action of a novel glue degrader AK59, which mediates interactions between HERC4, a HECT domain E3 ligase, and a novel substrate, STING. Molecular glue degraders and bi-functional PROTACS have attracted significant interest as a means to use small-molecule-derived changes in intracellular protein interactions for application to drug discovery ^37^. Recently, Liu et al. showed the usage of a known small molecule inhibitor against STING (C-170) combined with pomalidomide to achieve a PROTAC approach on STING degradation ^38^. While designing PROTACs from known inhibitors seemed to be a straightforward approach, specific glue degraders have significant advantages over PROTACs due to their size and therefore the easy conformity to Lipinski ‘s rule of five. Moreover, while CRBN-based degradation has been the focus of the field, this study is the first showing HERC4 as a potential E3 ligase for targeted protein degradation. This may be the first report of a compound directing a HECT domain containing E3 ligase to bring about targeted protein degradation.

Proteomics and genome-wide CRISPR/Cas9 screening techniques were then used to explore the mechanism of action of AK59 in an unbiased manner. By using a non-functional analog of AK59, QK50, both in proteomics as well as one-by-one interaction assays, it has been demonstrated that the chemical structure is unique and restrained in its ability for formation of the novel tripartite complex. While the ability of the compound to form this tripartite complex (STING, HERC4 and AK59) was validated using orthogonal assays, a number of questions remained. One of which is whether by creating a novel pocket, AK59 interacts at an interface between STING and HERC4 (so called proximal mechanism of action) or if by binding only one of the proteins, AK59 results in a conformational change which creates a novel PPI (so called distal mechanism of action) (see Figure 5F). Additionally, we managed to isolate the interaction site of STING to the cytosolic domain. In addition, it appears that AK59 was not competing with cGAMP binding but the specific degron sequence needed to be further investigated. Lastly, HERC4 binding site of AK59 still remains uncertain. To investigate these issues, compound structural-activity-relationship and protein structures needs to be further investigated with carefully selected assay readouts.

In dissecting AK59 ‘s function, the proteomics data showed at distinct cluster of proteins that are downregulated upon compound treatment (Supplementary Table S1). Among these proteins TMEM173 (STING) showed significant decrease in AK59 treatment sample compared to DMSO control and this significant decrease diminished in bortezomib-treated samples, indicating that the degradation of STING through AK59 is dependent on the proteasome (Supplementary Table S1). This observation was further supported by observation of K150 residue ubiquitination of STING in the presence of compound (Figure 2C). While we are mainly interested in the compound-mediated STING degradation, the proteomics data also shows a number of proteins additional to STING have been downregulated by AK59 treatment. While these proteins can be further investigated, the structure of AK59 could be optimized for specificity to STING recognition/degradation. Additionally, few proteins such as heme oxygenase 1 (HMOX1), members of heat shock protein Hsp70 family (HSPA1A, HSPA6) and members of heat shock protein Hsp40 family (DNAJA4, DNAJB4) are upregulated in the proteomics data. Elevated levels of these protein could indicate that the compound causes elevated ER stress, activation of proteolysis and protein quality control system ^39,40^.

This study reports the discovery of the first molecular glue degrader targeting STING, leading to inhibition of the cGAMP-activated IRF pathway. There has been numerous attempts to inhibit cGAS/STING pathway through targeting STING either via small molecule inhibitors, cGAMP derivatives or with PROTACs ^3,8,41^. To our knowledge, there has not previously been a report describing inhibitors, or degraders of STING, shown to be functional on SAVI mutants. While, the AK59 showed lower degradation of STING protein in the SAVI mutant lines, its novel mechanism as a molecular glue degrader using HERC4 as a novel E3 ligase opens new horizons for the regulation of STING in the context of autoimmune disorders.

## Material and Methods

### Cell culture

THP1 cells were cultured using Gibco RPMI-1640-Glutamax1 25mM Hepes medium (Life Technologies) supplemented with 10% FBS (PAA Laboratories), 1mM sodium pyruvate (Life Technologies) and 50 µM mercaptoethanol (Life Technologies). Cells were passaged every 4-5 days by dilution to 0.2-0.3 million cells/ml.

Dual-THP1 cells were cultured using Gibco RPMI-1640-Glutamax1 medium (Life Technologies) supplemented with 10% FBS (Life Technologies), 2mM L-glutamine (Life Technologies), 10mM Hepes (Life Technologies), 1mM sodium pyruvate (Life Technologies) and 50U/ml penicillin-streptomycin (Life Technologies). Cells were passaged to 0.2-0.3 million cells/ml every 4-5 days.

HEK293T or HEK293-JumpIN cells were cultured with Gibco DMEM with Glutamax (Life Technologies) supplemented with 10% FBS (Life Technologies), 10mM Hepes (Life Technologies) and 50U/ml penicillin-streptomycin (Life Technologies). Cells were passaged every 3-4 days, by diluting the cells 1 to 5, after releasing the attached cells using TrypLE (Life technologies).

For transfection studies, TransIT-LT1 transfection reagent (Mirus) was used according to the manufacturer’s protocol. For 6-well format, 1 million HEK293T cells were seeded one day before transfection. For each well, 2.5µg of DNA in 7.5µl of TransIT-LT1 transfection reagent diluted in 250µl optiMEM (Invitrogen). After 15 minutes of incubation of transfection reagent with DNA, mix was added dropwise on the cells. Then 48h after transfection, cells were treated with the compound of interest.

For lentiviral particle production, 1×10^7^ HEK293T cells were seeded on collagen coated T75 flasks (Thermo Fisher) and then 24 hours later, the cells were transfected with 1.84 µg of DNA of interest with 2.24 µg of ready-to-use lentiviral packaging plasmid mix (Cellecta) using Trans-IT transfection reagent (Mirus) according to the manufacturers ‘ protocol. Subsequently, the media was changed 24 hours after transfection and lentivirus-containing supernatants collected after 72 hours. Finally, lentivirus particles were filtered using a 0.45 µm filter (Millipore), before being concentrated 10-fold using the Lenti-X concentrator (Takara) according to the manufacturers ‘ protocol.

### Proteomics and analysis

THP1-Cas9 cells (2 ×10^6^ per well in a 6-well plate) were seeded, and treated, on the same day either with 0.1% (final) DMSO, 10 µM AK59 or 10 µM QK50 (0.1% DMSO final) for 16 hours. Cells were collected and washed with 1x DPBS (Gibco) after compound incubation. Before proteomics analysis of the samples the conditions were checked using western blot (for more details, see western blot) to confirm targeting effects on STING.

TMT-labeled peptides were generated with the iST-NHS kit (PreOmics) and TMT16plex reagent (Thermo Fisher Scientific). Equal amounts of labeled peptides were pooled and separated on a high pH fractionation system with a water-acetonitrile gradient containing 20 mM ammonium formate, pH 10 ^42^. Alternating rows of the resulting 72 fractions were pooled into 24 samples, dried and resuspended in water containing 0.1% formic acid.

The LC-MS analysis was carried out on an EASY-nLC 1200 system coupled to an Orbitrap Fusion Lumos Tribrid mass spectrometer (Thermo Fisher Scientific). Peptides were separated over 180 min with a water-acetonitrile gradient containing 0.1% formic acid on a 25 cm long Aurora Series UHPLC column (Ion Opticks) with 75 µm inner diameter. MS1 spectra were acquired at 120k resolution in the Orbitrap, MS2 spectra were acquired after CID activation in the ion trap and MS3 spectra were acquired after HCD activation with a synchronous precursor selection approach ^43^ using 8 notches and 50k resolution in the Orbitrap. LC-MS raw files were analyzed with Proteome Discoverer 2.4 (Thermo Fisher Scientific). Briefly, spectra were searched with Sequest HT against the *Homo sapiens* UniProt protein database and common contaminants (Sep 2019, 21494 entries). The database search criteria included: 10 ppm precursor mass tolerance, 0.6 Da fragment mass tolerance, maximum three missed cleavage sites, dynamic modification of 15.995 Da for methionines, static modifications of 113.084 Da for cysteines and 304.207 Da for peptide N-termini and lysines. The Percolator algorithm was applied to the Sequest HT results. The peptide false discovery rate was set to 1% and the protein false discovery rate was set to around 5%. TMT reporter ions of the MS3 spectra were integrated with a 20 ppm tolerance and the reporter ion intensities were used for quantification. All LC-MS raw files were deposited to the ProteomeXchange Consortium ^44^ via the PRIDE partner repository with the dataset identifier PXD00000. Protein relative quantification was performed using an in-house developed R (v.3.6) script. This analysis included multiple steps; global data normalization by equalizing the total reporter ion intensities across all channels, summation of reporter ion intensities per protein and channel, calculation of protein abundance log2 fold changes (L2FC) and testing for differential abundance using moderated t-statistics ^45^ where the resulting false rate discovery (FDR) p values (or q values) reflect the probability of detecting a given L2FC across sample conditions by chance alone. Raw data are available via ProteomeXchange with identifier PXD040291.

### Western blotting and co-immunoprecipitation

Cells were pelleted and washed with PBS (Gibco) before lysis. For western blot analysis, cell pellets were lysed with 5x extraction buffer (from co-immunoprecipitation kit, Thermo), diluted to 1x with 1xPBS (Gibco) and supplemented with cOmplete protease inhibitor (Roche) for 30 minutes on ice with pulse-vortexing every five minutes. To remove cell debris, lysate was spun at 15000 g at 4°C for 15 minutes. Supernatant, containing proteins were collected in a fresh tube and protein quantification was done using Pierce BCA assay (Thermo Fisher) according to the manufacturer’s protocol. Samples were prepared with NuPAGE LDS sample buffer (Invitrogen) and NuPAGE sample reducing agent (Invitrogen), boiled in 70°C for 10 minutes, loaded on pre-cast 10 well, 12 well or 15 well NuPAGE™ 4 to 12% Bis-Tris 1.5mm mini protein gel (Thermo Fisher) and run in 1x MES buffer (Invitrogen) for 45-50 minutes in 200V. Semi-dry transfer was done using Trans-blot turbo transfer (Biorad) with ready-made PVDF membrane-containing transfer packs according to the manufacturer’s protocol. Primary antibodies used in this study were all anti-human: STING (1:1000, CST, #13647 or 1:500, Thermo Fisher, MA526030), UBIQUITIN (1:500, CST, Cat#3936), HERC4 (1:500, abcam, Cat#ab856732), UBA6 (1:1000, CST, Cat#13386), UBA5 (1:500, abcam, Cat#ab177478), SMO (1:500, abcam, Cat#ab236465) and PGR (1:250, CST, Cat#8757). As loading control ACTIN (1:500, Sigma, Cat#A5441), TUBULIN (1:500, CST, Cat#2146) and VINCULIN (1:500, CST, Cat#13901) were used, as described in figure legends. HRP conjugated anti-mouse (1:2500, CST, Cat#7076) and anti-rabbit (1:2500, CST, Cat#7074) antibodies and Amersham ECL prime western blotting detection reagent (Cytiva Life Sciences) was used to detect protein levels. The ECL signal was visualized using Bio-rad ChemiDoc XRS+ and quantified using Image Lab program (Bio-rad).

Co-immunoprecipitation was performed using Dynabeads™ Co-Immunoprecipitation Kit (Thermo Fisher) according to the manufacturer’s protocol followed by western blot, which is described above. 50ul of HERC4 antibody (abcam, Cat# ab856732) conjugated with 10mg dynabeads and for each pulldown sample 1.5mg of antibody-conjugated dynabeads were used with 50ug of protein.

Co-immunoprecipitation for ubiquitin pulldown was performed using Ubiqapture-Q kit (Enzo Life Sciences) according to manufacturer’s protocols. To detect captured UBIQUITIN levels, instead of provided antibody from the kit, UBIQUITIN antibody from CST (mentioned above) was used.

### IRF pathway reporter assay

In order to measure IRF pathway activity, IRF-Lucia luciferase reporter system containing Dual-THP1-Cas9 cells (InvivoGen) were used. IRF pathway reporter assay was performed in clear bottom 96-well plates (TPP) where Dual-THP1-Cas9 cell line derivative cells were seeded at a cell density of 0.4 million cells/ well and stimulated with 30 µM cGAMP immediately. After three hours stimulation, cells were treated with the indicated compounds and the luciferase measurement was performed 16-hours after compound incubation. To detect the luciferase activity, QUANTI-Luc™ (Invivogen) was prepared according to the instructions on the data sheet. The reagent (50 µl) was added to a black with clear bottom 96-well plates (Greiner, Cat#655090) where the bottom of the plate was sealed (PerkinElmer, Cat#6005199). 20 µl of the Dual-THP1 cells from each condition pipetted on to the luminescence reagent in 96-well plate and luminescence was measured for 0.1 second with EnVision® Multimode plate reader. Fold increase in luminescence was calculated by dividing each read to its matching control group (DMSO treated) and results were plotted using Graphpad Prism 9.

### CRISPR genome-wide screening

For genome-wide CRISPR knockout screening, THP1 cells constitutively expressing Cas9 were generated by lentiviral delivery of Cas9 protein gene in pNGx-LV-c004 and selected with 5 µg/ml blasticidin S HCl (Thermo Fisher) as previously described ^46^. sgRNA library targeting 18,360 protein-coding genes with 5 sgRNA/gene, as previously described ^47^. The sgRNA library was packaged into lentiviral particles using HEK293T cells as previously described ^47,48^. Briefly, 2.1×10^7^ HEK293T cells were seeded into CellSTACK (Corning) cell culture chambers and transfected with the sgRNA library plasmid mix together with lentiviral packaging mix (Cellecta, containing psPAX2 and pMD2 plasmids that encode Gag/Pol and VSV-G, respectively) using Trans-IT transfection reagent (Mirus) 24 hours after seeding. Viral particles were harvested 72 hours post-transfection and quantified using LentiX qPCR kit (Clonetech).

THP1-Cas9 cells were expand for library transduction. On day 0, the cells were seeded and transfected to achieve a coverage of the library of at least 1000 cells/sgRNA with a multiplicity of infection (MOI) of 0.5. 5 µg/ml polybrene was used in transfection of THP1-Cas9 cells (Millipore). Transduced cells were selected with 4 µg/ml puromycin for three days and on the 4^th^ day, cells were analyzed for RFP expression using the FACS Aria for determining transduction efficiency. After sparing the day-4 samples, the rest of the library-transduced cells were seeded for the screen with two biological replicates per condition. At day 10, cells were treated with either DMSO or 10 µM AK59 and incubated for 16 hours. After compound incubation, cells were harvested, fixed, and stained for STING expression (described in detail below). Then the fraction of cells with the 25% high and 25% low level of staining for STING, in each treatment group the sorted samples were processed for genomic DNA isolation using the QIAamp DNA blood maxi kit (Qiagen) according to the manufacturer’s protocol. Genomic DNA was quantified using the Quant-iT PicoGreen assay (Invitrogen) according to manufacturer’s recommendations and proceeded with Illumina sequencing.

### Illumina sequencing of the library

The integrated gRNA sequences were PCR amplified using primers specific to the integrated lentiviral vector sequence and sequenced using the Illumina sequencing technology. Illumina library construction was performed as previously described ^47^. Briefly, a total of 96 µg of DNA per sample was split into 24 PCR reactions each with the volume of 100 µl, containing a final concentration of 0.5 µM of each of the following primers (Integrated DNA Technologies, 5644 5 ‘-AATGATACGGCGACCACCGAGATCTACACTCGATTTCTTGGCTTTATATATCTTGTGGAAAGGA-3 ‘ and INDEX 5 ‘-CAAGCAGAAGACGGCATACGAGATXXXXXXXXXXGTGACTGGAGTTCAGACGTGTGCTCTTCCGATC-3 ‘, where the Xs denote a 10 base PCR-sample specific barcode used for data demultiplexing following sequencing).

PCR samples were purified using 1.8x SPRI AMPure XL beads (Beckman Coulter) according to manufacturer’s recommended protocol and the qPCR products quantified using primers specific to the Illumina sequences using standard methods. Amplified libraries then pooled and sequenced with HiSeq 2500 instrument (Illumina) with 1x 30b reads, using a custom read 1 sequencing primer: 5645 (5 ‘-TCGATTTCTTGGCTTTATATATCTTGTGGAAAGGAC GAAACACCG-3 ‘), and a 1x 11b index read, using the standard Illumina indexing primer (5 ‘-GATCGGAAGAGCACACGTCTGAACTCCAGTCAC-3 ‘), according to the manufacturer’s recommendations.

### Analysis of the CRISPR screen

Sequencing analysis was performed as previously described ^47^. In short, raw sequencing reads were converted to FASTQ format using bcl2fastq2 (version 2.17.1.14, retrieved from http://support.illumina.com/downloads/bcl2fastq-conversion-software-v217.html); trimmed to the guide sequence with the fastx-toolkit (version 0.0.13, retrieved from http://hannonlab.cshl.edu/fastx_toolkit/index.html) and aligned to the sgRNA sequences in the library using bowtie ^49^ with no mismatches allowed. Differential expression of sgRNAs was calculated using DESeq2 ^50^ and gene-level results were obtained using the redundant siRNA activity (RSA) algorithm ^51^. In RSA, the rank distribution of individual sgRNAs is examined to calculate a hypergeometric enrichment score for the concerted action of each gene ‘s set of guides. This results in a gene-level p-value for significance, and we also use the lower (Q1) or upper (Q3) quartile of the guide ‘s fold changes to represent effect size at the gene level.

Due to the pooled analysis approach in RSA algorithm, range of RSA and Q values in each comparison is in the similar magnitude which allow us to do further comparisons. In order to rank the hits, RSA and Q values from two different comparisons (in this case DMSO high vs. low STING and AK59 high vs. low STING) were taken as point coordinates. Between two coordinates (aka RSA and Q values of each comparison) the Euclidean distance were calculated as the magnitude of the vector and the direction of the vector was assigned by the increase/decrease of RSA and Q values between two points (Bioconductor 4.0.2). Then, hits were ranked according to the magnitude and the direction of their vector.

STRING analysis was performed only between 3 selected hits (HERC4, UBA5 and UBA6) and STING to represent the relation of the genes with each other (STRING version 11.5) ^34^.

### STING quantification by flow cytometry

THP1-Cas9 cells were seeded at a density of 1×10^6^ cells/ml in 24 well plates with the media containing the indicated compound. Compound incubation time varied between 5-16 hours and bortezomib treatments were always 1 hour prior to any additional compound treatment. Cells were always seeded together with the initial treatment. After the compound incubation, cells were collected and fixed using 2.5% paraformaldehyde (stock 32%; # 15714-S; Electron Microscopy Sciences) at 37°C for 10 minutes. Cells were then washed using FACS Wash Buffer containing 1x D-PBS + 0.5% FBS + 2mM EDTA and permeabilized at room temperature for 20 minutes using 100µL of Perm/Wash I (BD # 557885), diluted 1:10 with 1x D-PBS. Cell washing was performed again and then samples were stained with 150µL/sample anti STING Alexa488; 1:200 (anti TMEM173; Abcam # ab198950) diluted in Robosep Buffer, which contained PBS + 2.0% FBS + 1mM EDTA (Stemcell; # 20104) at 4°C for 1-2 hours. After antibody incubation, cells were washed with Wash Buffer three times then cell pellets were resuspended in FACS Wash Buffer and flow cytometry acquisition on Fortessa was then performed. Analysis was performed using FlowJo software (version 10.6.1)

### Individual CRISPR knockouts and TIDE analysis

THP1-Cas9, Dual-THP1-Cas9 ^24^ and HEK293-JumpIN-Cas9 cells were generated by lentiviral delivery of Cas9 protein gene in pNGx-LV-c004 ^46^ and selected with 5 µg/ml blasticidin S HCl (Thermo Fisher). Individual knockouts were generated by lentiviral delivery of sgRNAs in the pNGx-LV-g003 backbone. SgRNA transduced cells were selected with 3.5 µg/ml puromycin (Thermo Fisher). sgRNA sequences targeting the gene of interests are listed below. Knockout efficiency checked by both Tracking of Indels by Decomposition (TIDE) analysis ^35^ and western blot.

To check the rate of Indel formation at the CRISPR cut site, TIDE analysis ^35^ was used as previously described ^52^. Briefly, genomic DNA was extracted from approximately 1 million cells per condition using DNA extraction kit (Qiagen) according to the manufacturer’s recommendations. DNA concentration is measured with Nanodrop (Thermo Fisher), and PCR reaction was performed with 2x Phusion polymerase master mix (Thermo Fisher), 5% DMSO, 100ng of DNA and final concentration of 1 µM forward and reverse primer. PCR primers for each gene listed below. Amplification run was initial denaturation in 98°C 30 seconds, 30 cycles of 98°C 5 seconds, 61°C 10 seconds, 72°C 15 second and final extension of 72°C for 2 minutes. PCR samples were cleaned up using PCR and Gel extraction kit (Qiagen) according to the manufacturer’s protocol and Sanger sequenced.

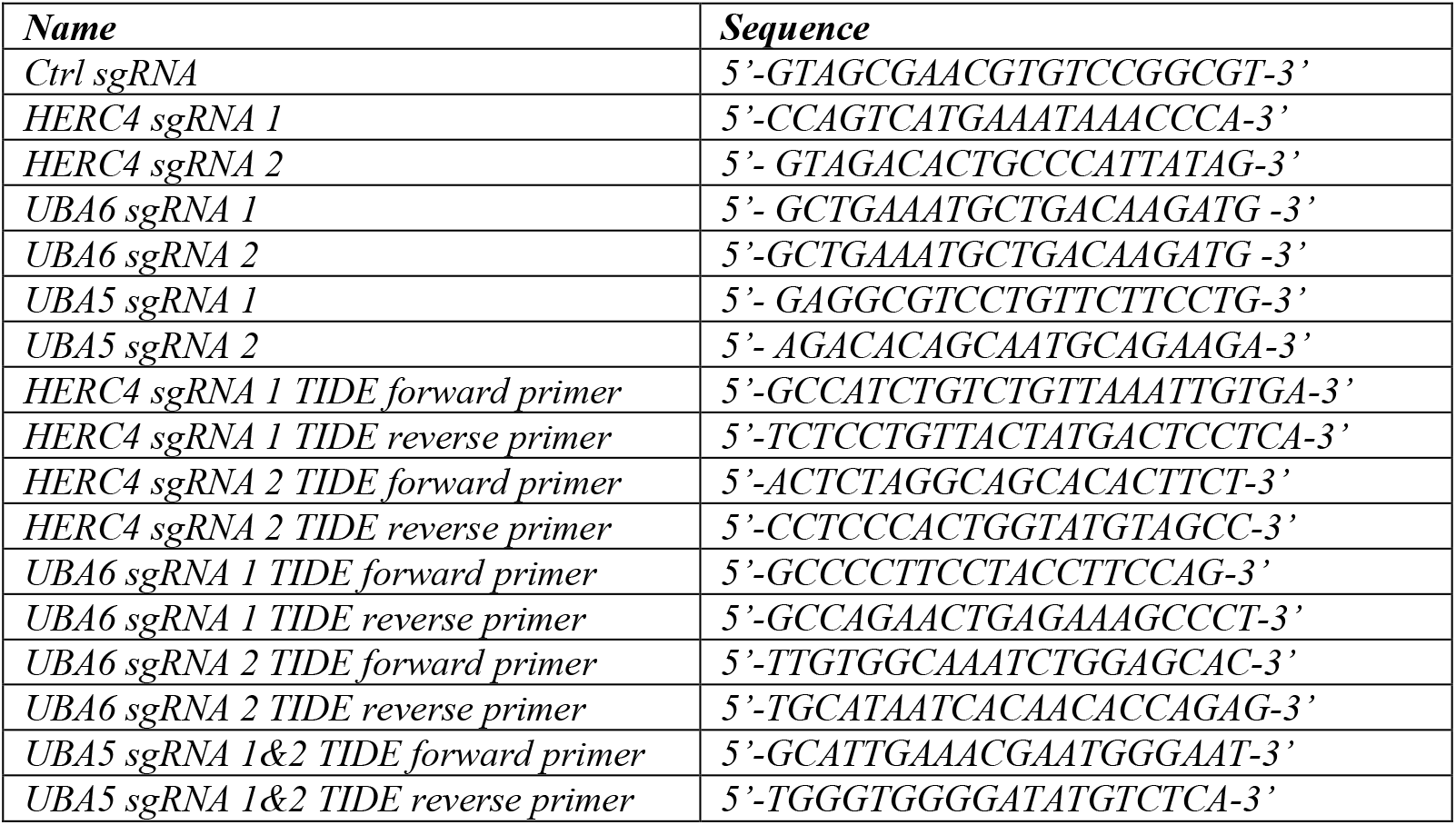

### STING-GFP construct design and tracking with live cell imaging

Cytosolic domains for STING (consisting of either the C-terminal 141-341aa or 155-341aa) were cloned into the pcDNA3.1(+) (Thermo Fisher) vector backbone with a C terminal GFP tagging. HEK293T cells were seeded on 6-well plates (TPP) with a cell density of 1 million cells/well. One day after seeding, cells were transfected with the STING-GFP expression constructs using the Trans-IT transfection reagent (Mirus). After transfection, cells were placed into an Incucyte® (Sartorius) live cell imager. 48-hour after transfection, cells were treated with either DMSO or 10 µM AK59. Cells were treated with compound for 16 hours. Live cell analysis was performed using Incucyte software where 10x phase contrast and GFP images were taken of each well every 30-45 minutes; 16 frames were set on each well for even quantification and data is normalized to start of the treatment. For the HERC4 knockout and Ctrl sgRNA comparison, all the wells were treated with the AK59 and the GFP signal difference between STING^155-341^ and STING^141-3431^ was measured.

### Nanobit complementation assay

Full length human HERC4 (CCDS41533) was cloned into the pFN35K SmBiT TK-neo Flexi® Vector, N-terminal small bit (SmBiT) tagged construct (Promega), with the native HSV-TK promoter swapped out for a CMV promoter. Cytosolic domain of human STING (141-341aa) was cloned in pFC34K LgBiT TK-neo Flexi® Vector, C-terminal large bit (LgBiT) tagged construct (Promega) which is transcribed under control of the HSV-TK promoter.

Next, 4×10^4^ HEK293T cells were seeded into of 96-well plates (TPP). On the same day, cells were transiently transfected with both of the constructs using Trans-IT transfection reagent (Mirus) according to the manufacturer’s recommended protocol. Then 48 hours after transfection, cells were treated indicated doses of with either DMSO, AK59 or YJX209. Luminescence activity was the measured 30 minutes, 5 hours, 8 hours and 12 hours after compound incubation. For the dose response, all samples were measured at 16-hour post compound treatment. For the cGAMP competition with AK59, cGAMP stimulation was initiated by the addition of 30 µM of cGAMP three hours prior to the 16-hour compound treatment. To detect the luciferase activity, Nano-Glo® luciferase assay system (Promega) was used according to manufacturer’s protocol and luminescence was measured for 0.1 second with EnVision® Multimode plate reader. Fold increase in luminescence was calculated by dividing each read to its matching control group (DMSO treated) and results were plotted using Graphpad Prism 9.

### Statistics

Statistical analysis of the indicated data was performed using Graphpad Prism 9. Data points were represented as the mean ± standard deviation (SD). Appropriate statistical test for each data indicated in the figure legends.

## Acknowledgments

This work was supported by Discovery Postdoc Program in Novartis Institutes of BioMedical Research (NIBR) Postdoc Office.

## Contributions

M.M., I.S., A.H., C.N.P. and D.G. designed the research. M.M., I.S., A.I.M., B.G., F.F., D.B., S.K., S.R., D.B., D.J., T.T., R.M., S.A. and A.R. performed research. M.M., N.P., E.A. and A.H. analyzed data. M.M., F.N., A.H., J.R.H., C.N.P. and D.G. wrote and edited paper.

## Supplementary Materials

Supplementary Table S1. Proteomics results from either AK59 or QK50 treated THP1 cells.

Supplementary Table S2. CRISPR/Cas9 knockout screen RSA to Q values of hits.

**Supplementary Figure 1.**
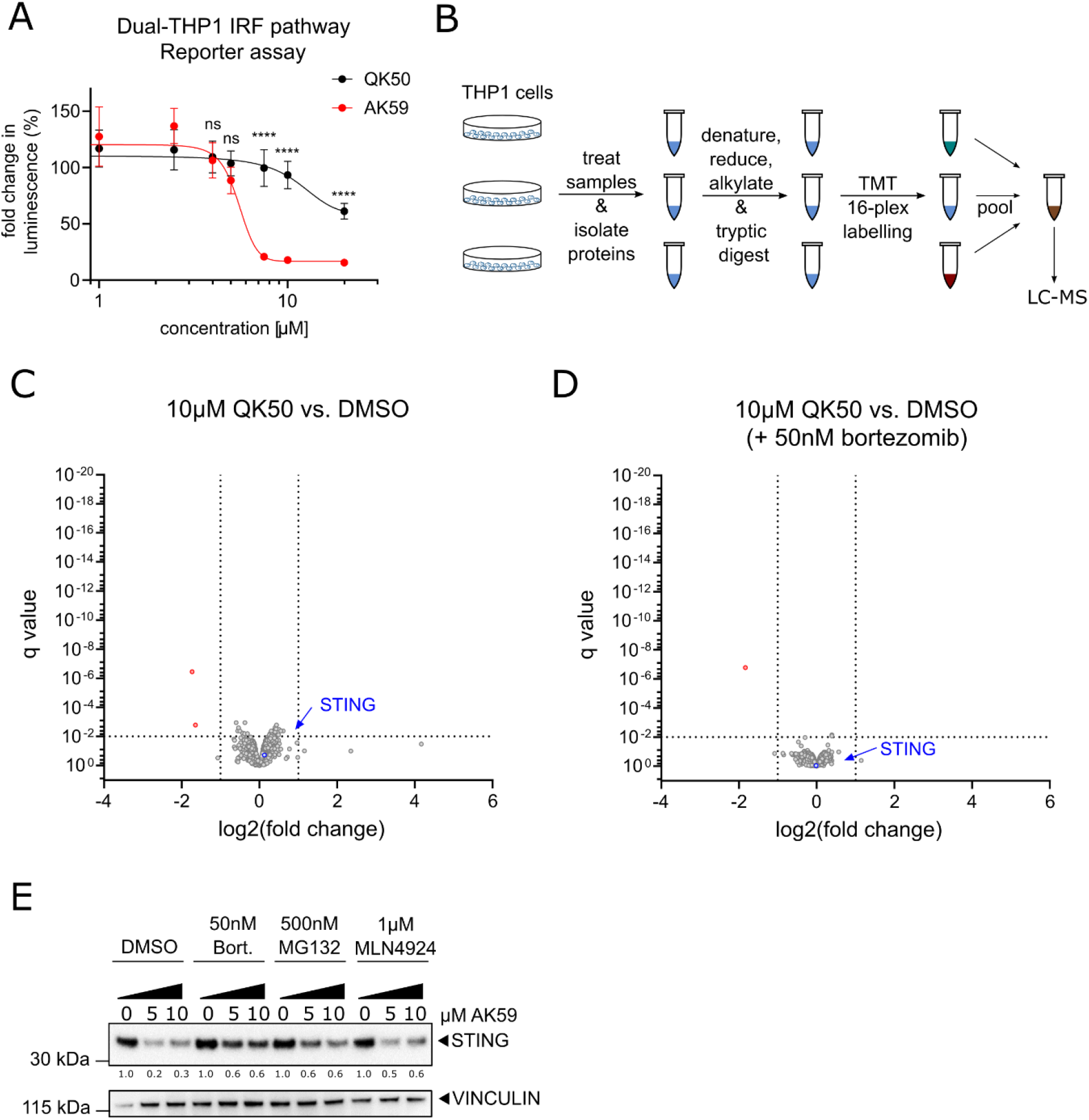
Compound characterization of AK59 and QK50. **(a)** IRF pathway reporter assay on wild type Dual-THP1 cells treated with either AK59 or QK50 for 16 hours. Cells were stimulated with 30 µM cGAMP 3 hours prior to compound incubation. Luminescence reads were normalized to the control unstimulated sample for each cell line. Data plotted as mean ± SD of three individual biological replicates in Graphpad Prism (Version 9). Statistical significance was calculated using two-way ANOVA followed by Šidák ‘s correction. **(b)** Schematic representation of proteomics on THP1 cells treated with either DMSO, AK59 or QK50. **(c-d)** Volcano plots of the proteomics analysis of THP1 cells treated for 16 hours with 10 µM QK50 and with or without 50nM of bortezomib treatment compared to matching DMSO control samples. Significantly altered protein abundances are shown with a log2 fold change < - 1 or > 1 and a q value of less than 0.01. **(e)** Western blot representing the changes in STING expression upon AK59 and/or proteasomal inhibition (bortezomib or MG132) or neddylation inhibition (MLN4924) in THP1 cells. VINCULIN used as a loading control. The STING expression quantified by first normalizing to its respective loading control and ratio to DMSO control.

**Supplementary Figure 2.**
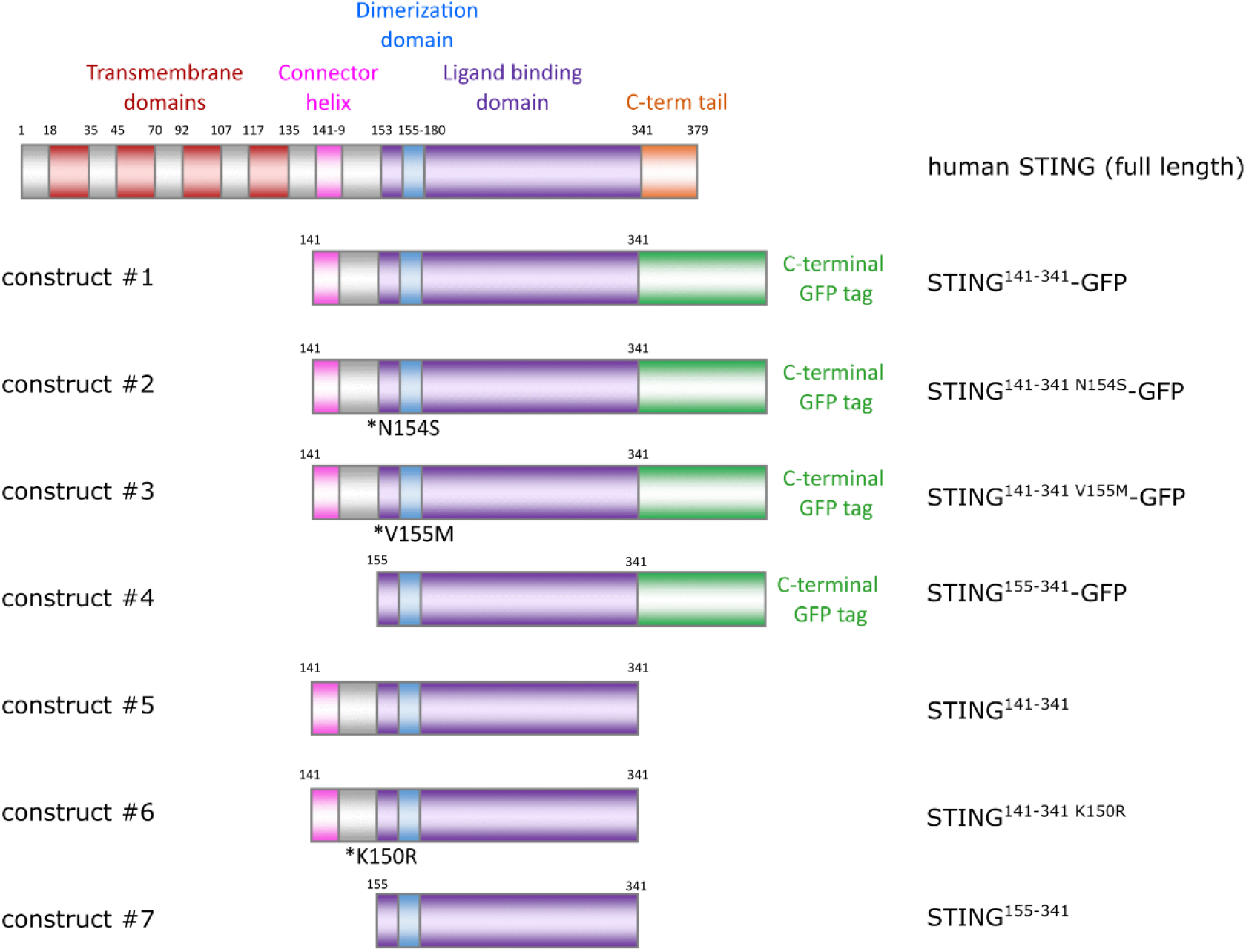
STING constructs. Schematic representation of the STING constructs that are transiently expressed in HEK293T cells.

**Supplementary Figure 3.**
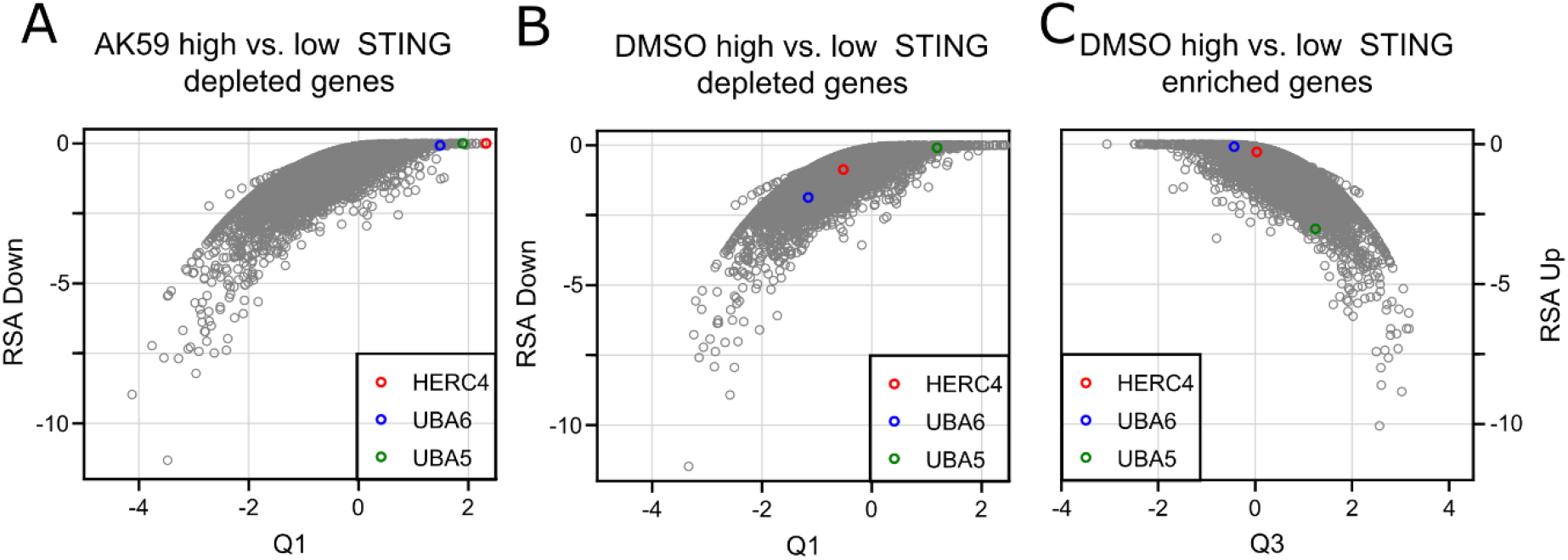
CRISRP/Cas9 genome-wide screening results. **(a)** RSA down to Q1 values from CRISPR/Cas9 screen of 10 µM AK59 treated THP1-Cas9 cells represented in a mustache plot. The comparison represented in the blot is high STING expression to low STING expression in the AK59 treatment group. Each dot represents a gene from the CRISPR/Cas9 library. **(b)** RSA up to Q3 values from CRISPR/Cas9 screen of DMSO treated THP1-Cas9 cells represented in a mustache plot. The comparison represented in the blot is high STING expression to low STING expression in the DMSO treated group. Each dot represents a gene from the CRISPR/Cas9 library. **(c)** RSA down to Q1 values from CRISPR/Cas9 screen of DMSO treated THP1-Cas9 cells represented in a mustache plot. The comparison represented in the blot is high STING expression to low STING expression in the DMSO treated group. Each dot represents a gene from the CRISPR/Cas9 library.

**Supplementary Figure 4.**
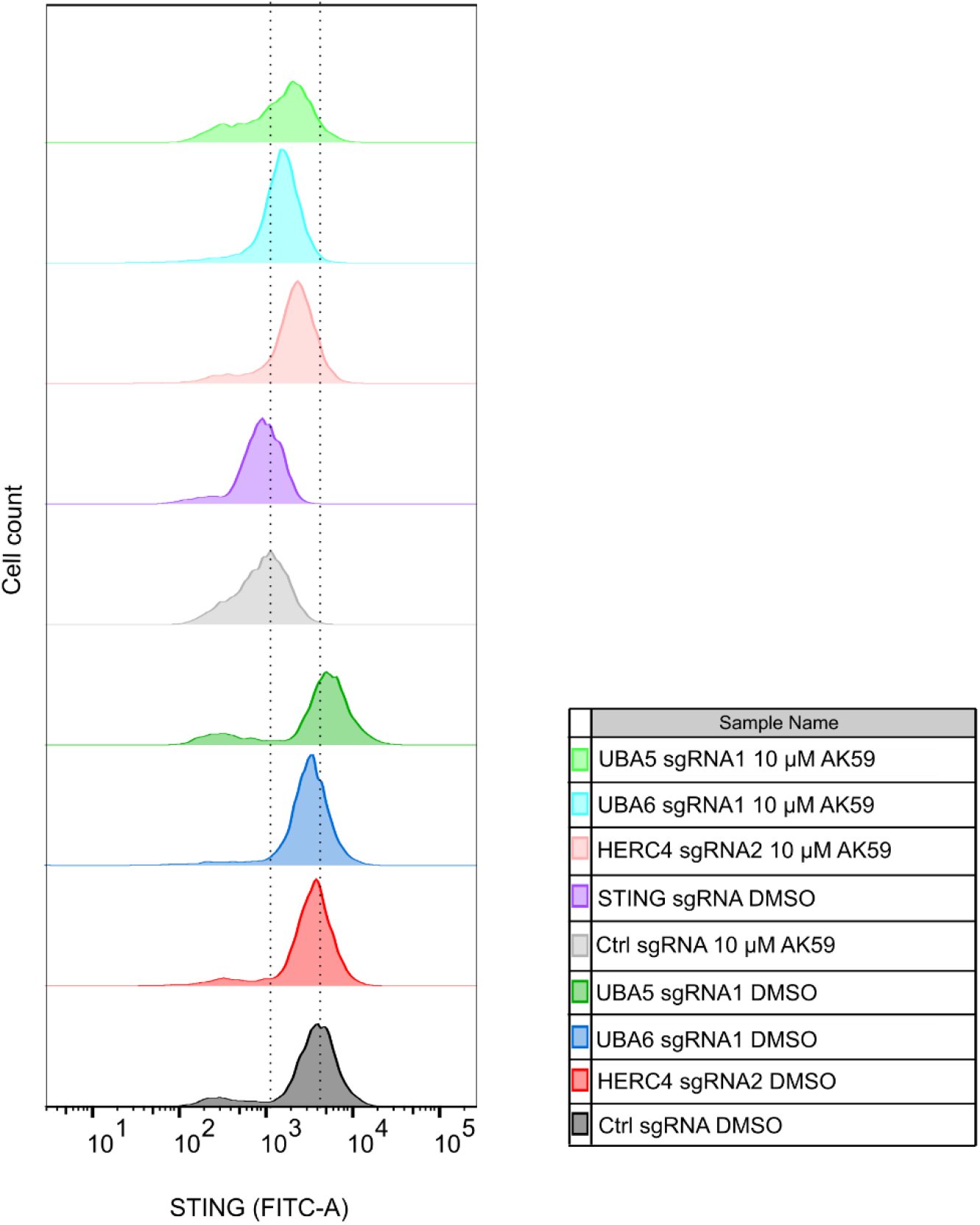
HERC4, UBA5 and UBA6 are validated as genes responsible in AK59 activity on STING expression. FACS analysis of STING expression on Ctrl sgRNA, HERC4 sgRNA2, UBA6 sgRNA1 or UBA5 sgRNA1 transduces THP1-Cas9 cells that are treated either with DMSO or 10 µM AK59. STING knockout line was taken along as a negative control (purple). Experiment treated at least in three biological replicates and representative FACS reads plotted using FlowJo (Version 10.6.1).

**Supplementary Figure 5.**
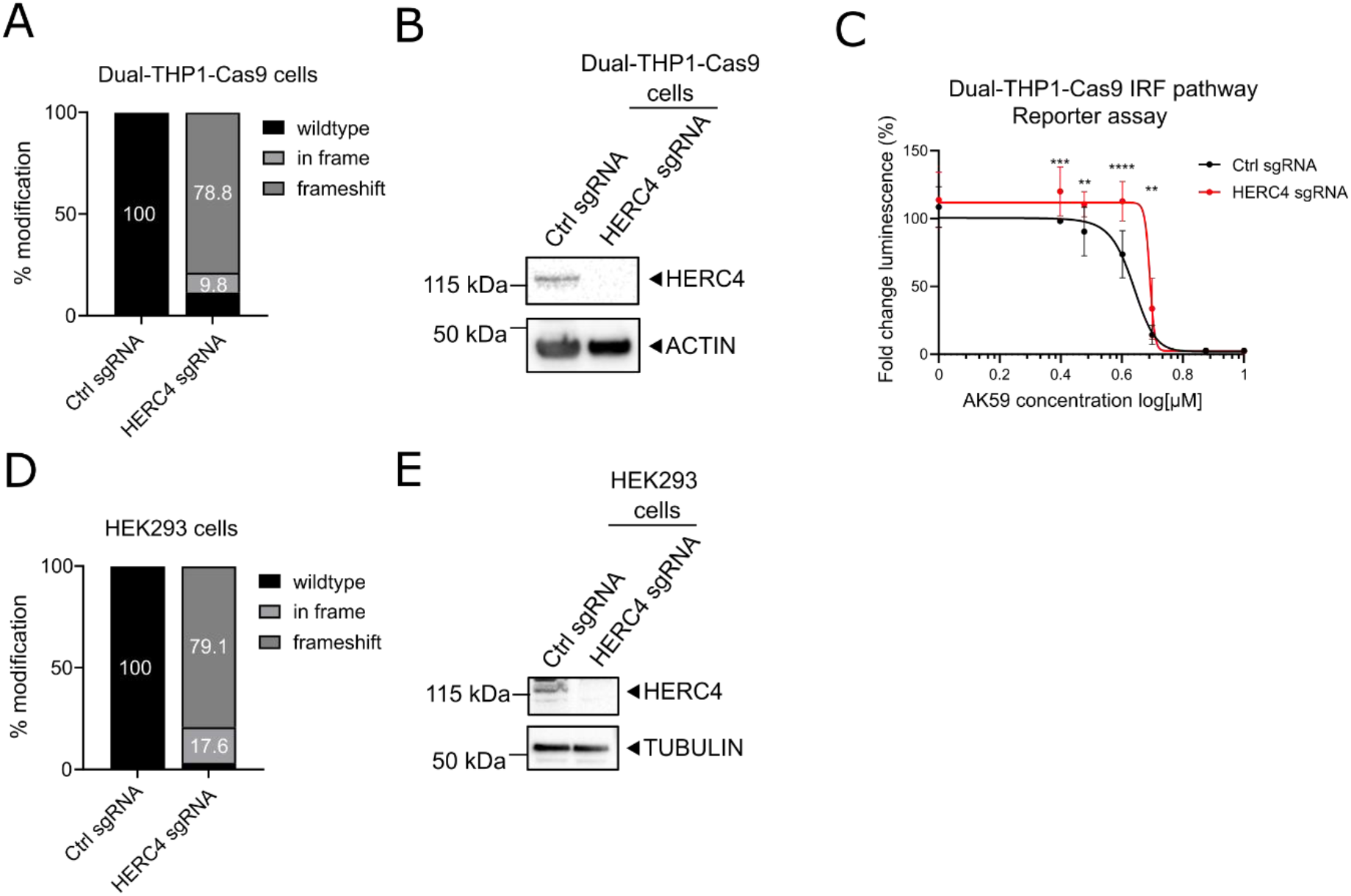
Confirmation HERC4 knockouts in HEK293T and Dual-THP1 cells. **(a)** TIDE analysis from either Ctrl sgRNA or HERC4 sgRNA1 transduced Dual-THP1-Cas9 cells. Modification rates were checked at the third passage after the initial transduction. Undefined sequencing reads were excluded. **(b)** Western blot to detect HERC4 protein levels on either Ctrl sgRNA or HERC4 sgRNA1 transduced Dual-THP1-Cas9 cells. Proteins from each cell line collected at the third passage after the initial transduction. **(c)** IRF pathway reporter assay on wild type or HERC4 knockout Dual-THP1-Cas9 cells. Cells were stimulated with 30 µM cGAMP 3 hours prior to 16 hours AK59 incubation. Luminescence reads were normalized to the control unstimulated sample for each cell line. Data plotted as mean ± SD of three individual biological replicates in Graphpad Prism (Version 9). Statistical significance was calculated using two-way ANOVA followed by Šidák ‘s correction. **(d)** TIDE analysis from either Ctrl sgRNA or HERC4 sgRNA1 transduced HEK293-JumpIN-Cas9 cells. Modification rates were check at the third passage after the initial transduction. Not defined sequencing reads were excluded. **(e)** Western blot to detect HERC4 protein levels on either Ctrl sgRNA or HERC4 sgRNA1 transduced HEK293-JumpIN-Cas9 cells. Proteins from each cell line collected at the third passage after the initial transduction.

**Supplementary Figure 6.**
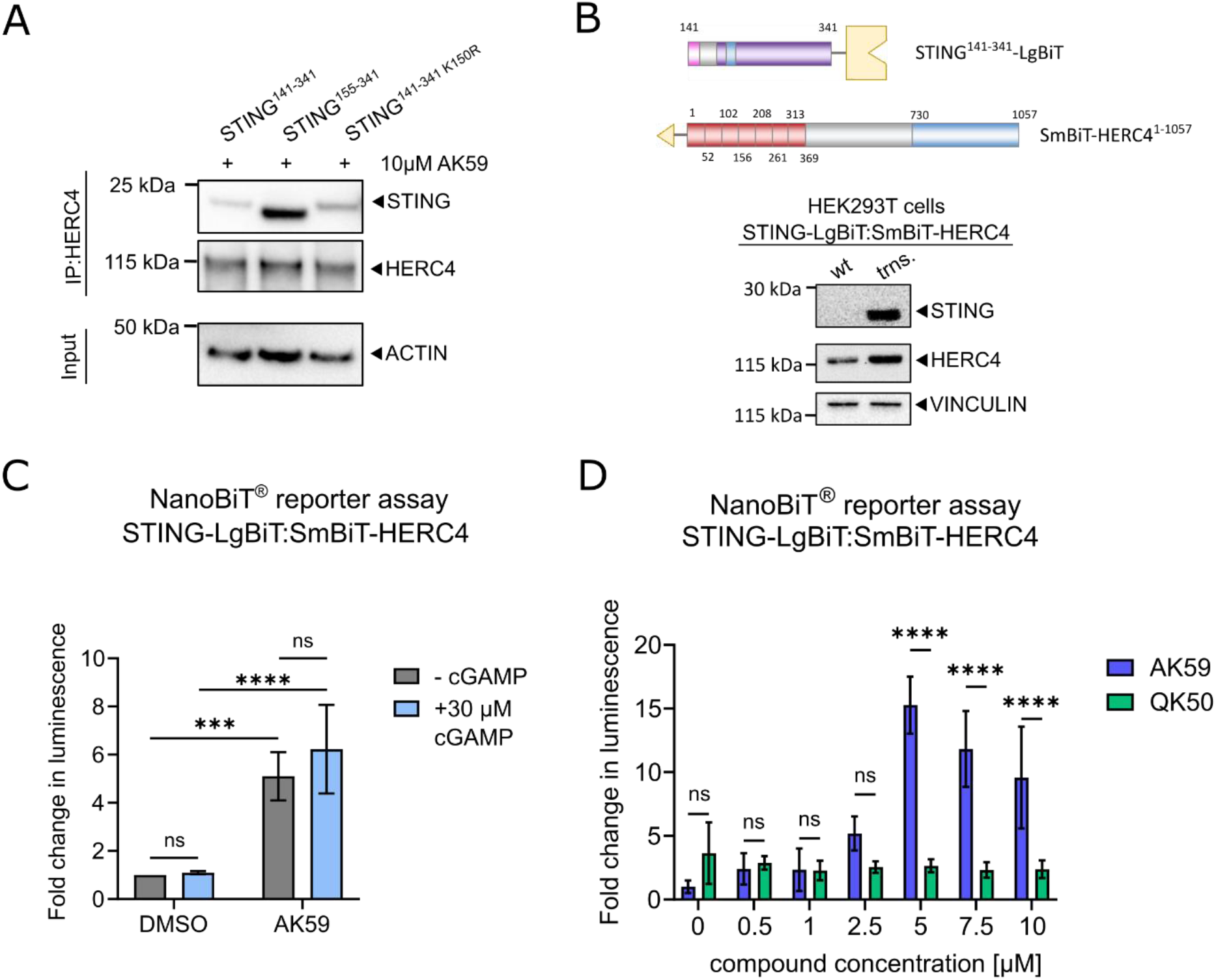
Interaction of HERC4 and STING in the presence of AK59 treatment. **(a)** HERC4 pulldown followed by western blot on HEK293T cells transfected with the indicated STING expression constructs and treated with 10 µM AK59. ACTIN was used as a loading and pulldown control. **(b)** Schematic representation of the Nanobit constructs that were transiently expressed in HEK293T cells. Western blot showing the expression of STING-LgBiT and SmBiT-HERC4 constructs in HEK293T cells. Comparison of wildtype (wt.) and transfected (trns.) HEK293T cells. VINCULIN was used as a loading control. **(c)** Nanobit complementation assay on STING-LgBiT and SmBiT-HERC4 expressing HEK293T cells in the presence of AK59 and/or cGAMP. 30 µM of cGAMP stimulation was done 3 hours prior to indicated compound treatments. DMSO were used as a negative control and luminescence reads were normalized to corresponding DMSO control. Three biological replicates were plotted as mean ± SD using in Graphpad Prism (Version 9). Statistical significance was calculated using two-way ANOVA followed by Šidák ‘s correction. **(d)** Nanobit complementation assay on STING-LgBiT and SmBiT-HERC4 expressing HEK293T cells in the presence of increasing concentration of either AK59 or QK50. Biological replicates were plotted as mean ± SD using in Graphpad Prism (Version 9). Statistical significance was calculated using two-way ANOVA followed by Šidák ‘s correction.

